# Integrative genetic and structural analyses reveal nitric oxide-driven regulation of ERFVII stability in Arabidopsis under low O_2_

**DOI:** 10.64898/2026.07.28.741153

**Authors:** Ashim Kumar Das, Hammad Ismail, Da-Sol Lee, Madiha Hameed, Sang-Mo Kang, Mohammad Golam Mostofa, Byung-Wook Yun

## Abstract

Nitric oxide (NO) is a key signaling molecule that regulates diverse physiological responses, including adaptation to hypoxia in plants. Although stabilization of group-VII ETHYLENE RESPONSE FACTOR (ERFVII) transcription factors under low O_2_ is known to be facilitated by the N-dragon pathway, whether NO directly mediates N-terminal cysteine (Cys) oxidation to regulate ERFVII stability remains unresolved. Here, we combined genetic, computational, and structural approaches to investigate NO’s role in proteasomal degradation of ERFVII during flooding stress. Arabidopsis mutants with elevated S-Nitrosoglutathione (GSNO) levels compromised ERFVII activation during dark submergence but increased expression of genes encoding N-degron pathway enzymes. Although direct detection of *in vivo* NO-mediated S-nitrosylated proteins remains technically challenging, the GPS-SNO 1.0 tool predicted the conserved N-terminal second Cys residue as a high-confidence S-nitrosylation site. Structural modeling using the Schrödinger Suite 2024-4 further revealed that conversion of the Cys thiol (Cys-SH) to S-nitrosothiol (Cys-SNO) induced notable conformational changes in ERFVII TFs. Molecular docking further demonstrated that Cys-SNO-modified ERFVII peptides exhibited stronger binding affinities and altered interaction networks with N-degron pathway enzymes, supporting a role for NO-mediated structural remodeling in ERFVII degradation. Elevated GSNO also disrupted energy balance efficiency for overcoming O_2_ deficiency, altered the expression of sugar starvation-responsive genes, and impaired ATP binding capacity of Cys-SNO ERFVII proteins, as evidenced by docking and molecular dynamics simulations. Collectively, these results support a model in which NO-mediated modification of the conserved N-terminal Cys promotes ERFVII degradation, thereby linking NO signaling to hypoxia-responsive transcriptional regulation and metabolic adaptation during flooding stress.

## Introduction

Flooding is a major environmental constraint that limits plant growth and productivity by restricting oxygen (O_2_) availability. To survive hypoxia, plants rapidly reprogram their metabolism, growth, and gene expression through an intricate signaling network in which ethylene and nitric oxide (NO) play central roles ^1^. Ethylene is a gaseous plant hormone that regulates numerous developmental and stress-responsive processes ^2^. Under flooding or O_2_-deficient conditions, reduced gas diffusion leads to ethylene accumulation in plant tissues, where it initiates adaptive responses to O_2_ deprivation ^3^. A central component of ethylene-mediated O_2_-sensing is the group VII ETHYLENE RESPONSE FACTOR (ERFVII) family of APETALA2/ERF-type transcription factors (TFs) ^4,5^, which activate hypoxia-responsive genes to sustain energy production, conserve metabolic resources, protect subcellular components, and mitigate cellular damage under anaerobic conditions ^6^.

The biological importance of ERFVII transcription factors has been demonstrated across plant species. In rice (*Oryza sativa*), the ERFVII TF SUBMERGENCE 1A-1 (SUB1A-1) conferred high tolerance to prolonged submergence through quiescence mechanisms ^7,8^, whereas closely related alleles, including SUB1A-2, SUB1B, and SUB1C, showed an intolerant phenotype to submergence ^8^. On the other hand, deepwater rice uses an escape strategy by the ERFVII TF SNORKEL1 and SNORKEL2, which promote gibberellin- and ethylene-dependent internode elongation during submergence ^9^. In *Arabidopsis thaliana*, five ERFVII members-*RELATED TO APETALA2.12* (*RAP2.12*), *RAP2.2*, *RAP2.3*, *HYPOXIA-RESPONSIVE ERFs* (*HRE1*), and *HRE2*-coordinate hypoxia adaptation by activating anaerobic genes by binding to the hypoxia-responsive promoter element (HRPE) ^10^.

The abundance of most ERFVII proteins is regulated by the O_2_-dependent Arg/Cys branch of the N-degron pathway, a proteolytic mechanism in which protein stability is determined by the identity and modification state of the amino-terminal residue ^11,12^. In the group VII ERF family, the Nt-Cysteine (Cys) residue, exposed upon cleavage of methionine by MAP (Methionine aminopeptidase) enzymes, is susceptible to oxidation. In the presence of O_2_, Cys oxidation (*Cys) takes place by plant Cys oxidases (PCOs), which convert Cys into Cys-sulfinic acid, while O_2_ deficiency inhibits PCOs in facilitating the oxidation process ^13^. The oxidized residue is subsequently arginylated by arginyl transferases (ATEs), enabling recognition by the E3 ubiquitin ligase PROTEOLYSIS6 (PRT6) for proteasomal degradation ^6,14^. Under hypoxia, reduced PCO activity stabilizes ERFVII proteins, allowing activation of O_2_-deficiency responses. Interestingly, SUB1A-1 largely escapes this canonical degradation pathway despite possessing the conserved N-degron motif, suggesting additional regulatory mechanisms modulate ERFVII stability ^15^.

NO has emerged as another critical regulator of ERFVII turnover, yet its precise role remains controversial. Besides O_2_, Nt-Cys is reported to be oxidized non-enzymatically by NO ^16^. Early studies proposed that NO promotes degradation of ERFVII proteins through oxidation of the Nt-Cys, while ethylene-induced scavenging of NO by PHYTOGLOBIN1 (PGB1) enhances RAP2.12 stability under hypoxia ^17–19^. However, subsequent biochemical studies questioned whether NO directly oxidizes the N-terminal Cys, suggesting instead that NO-dependent degradation may occur through non-enzymatic reactions or mechanisms independent of prior Cys oxidation ^13,20,21^. Consequently, the molecular relationship between NO signaling and the Arg/Cys branch of the N-degron pathway remains unresolved.

A major mechanism of NO signaling is S-nitrosylation, a reversible post-translational modification in which NO covalently modifies Cys thiols to form S-nitrosothiol (SNO) ^22,23^. Cellular S-nitrosylation levels are primarily regulated by S-nitrosoglutathione reductase (GSNOR), which controls the abundance of the NO reservoir S-nitrosoglutathione (GSNO) ^24^. S-nitrosylation is increasingly recognized as a regulator of protein stability through the ubiquitin–proteasome system. In Arabidopsis, S-nitrosylation of ASCORBATE PEROXIDASE 1 (APX1) and ABA INSENSITIVE 5 (ABI5) causes their degradation through the ubiquitin–proteasome pathway, whereas it protects NON-EXPRESSOR OF PATHOGENESIS-RELATED GENES1 (NPR1) from ubiquitination ^25–27^. Moreover, a loss-of-function mutation in *gsnor1-3* exhibited reduced germination under O_2_-deficient conditions, and GSNOR1 degradation occurred via autophagy instead of ubiquitylation ^28^. Despite these advances, whether S-nitrosylation directly modifies the conserved N-terminal Cys of ERFVII proteins and thereby influences their degradation through the N-degron pathway has not been investigated. Here, we tested the hypothesis that NO regulates ERFVII stability through S-nitrosylation of the conserved Nt-Cys, thereby modulating recognition by the N-degron machinery during hypoxia. We combined genetic analyses of *Arabidopsis* GSNOR1 loss- and gain-of-function mutants with structural modeling, molecular docking, and molecular dynamics simulations to examine how NO-mediated redox modification influences ERFVII conformation, interactions with N-degron pathway components, and hypoxia-responsive gene expression. We further evaluated the consequences of altered GSNO homeostasis for submergence tolerance, energy metabolism, and sugar-starvation responses. Together, this study provides mechanistic insights into how NO-dependent redox regulation may integrate with O_2_ sensing to control ERFVII stability and flooding adaptation in plants.

## Results

### Elevated GSNO enhances sensitivity to submergence and potentiated proteolytic degradation

Based on previous reports, we know that group VII ERF TFs are less stable in the presence of NO, which regulates seed germination, limits hypocotyl elongation, and promotes stomatal closure ^17^. NO disproportionately controls ethylene levels under O_2_-deficient conditions ^19^. Although non-enzymatic NO responses are believed to regulate Nt-Cys oxidation, yet not fully understood, nor are there sufficient reports of submergence tolerance in NO mutants. In this study, we investigated NO responses to submergence using loss- and gain-of-function mutations in GSNOR1, namely *gsnor1-3* and *gsnor1-1*, respectively, which have been shown to regulate cellular SNO levels and likely control global S-nitrosylation ^22^. As shown in Figure 1A, four-week rosette plants of *gsnor1-3* and *gsnor1-1*, along with Col-0, were subjected to dark submergence for 5 days. Phenotypic evaluation showed that elevated GSNO levels in *gnsor1-3* increase sensitivity to low O_2_, as indicated by severe leaf rotting at day 5 (Figure 1A). In contrast, the low O_2_ sensitivity was largely compromised in *gsnor1-1* and Col-0 plants. Consistent with this result, the percentage of damaged leaves also showed less damage in *gsnor1-1* and Col-0, compared to *gsnor1-3*, over three consecutive days of dark submergence (Figure 1B). Col-0 and *gsnor1-1* plants outperformed *gsnor1-3* in maintaining photosynthetic pigment levels during dark submergence (Figure 1C–E). Moreover, we found that *gsnor1-1* plants had lower H_2_O_2_ levels but higher GSH levels, whereas *gsnor1-3* plants had lower GSH levels (Supplementary Figure 1). Collectively, these results indicate that GSNO negatively regulates the O_2_-deficient environments.

**Figure 1.**
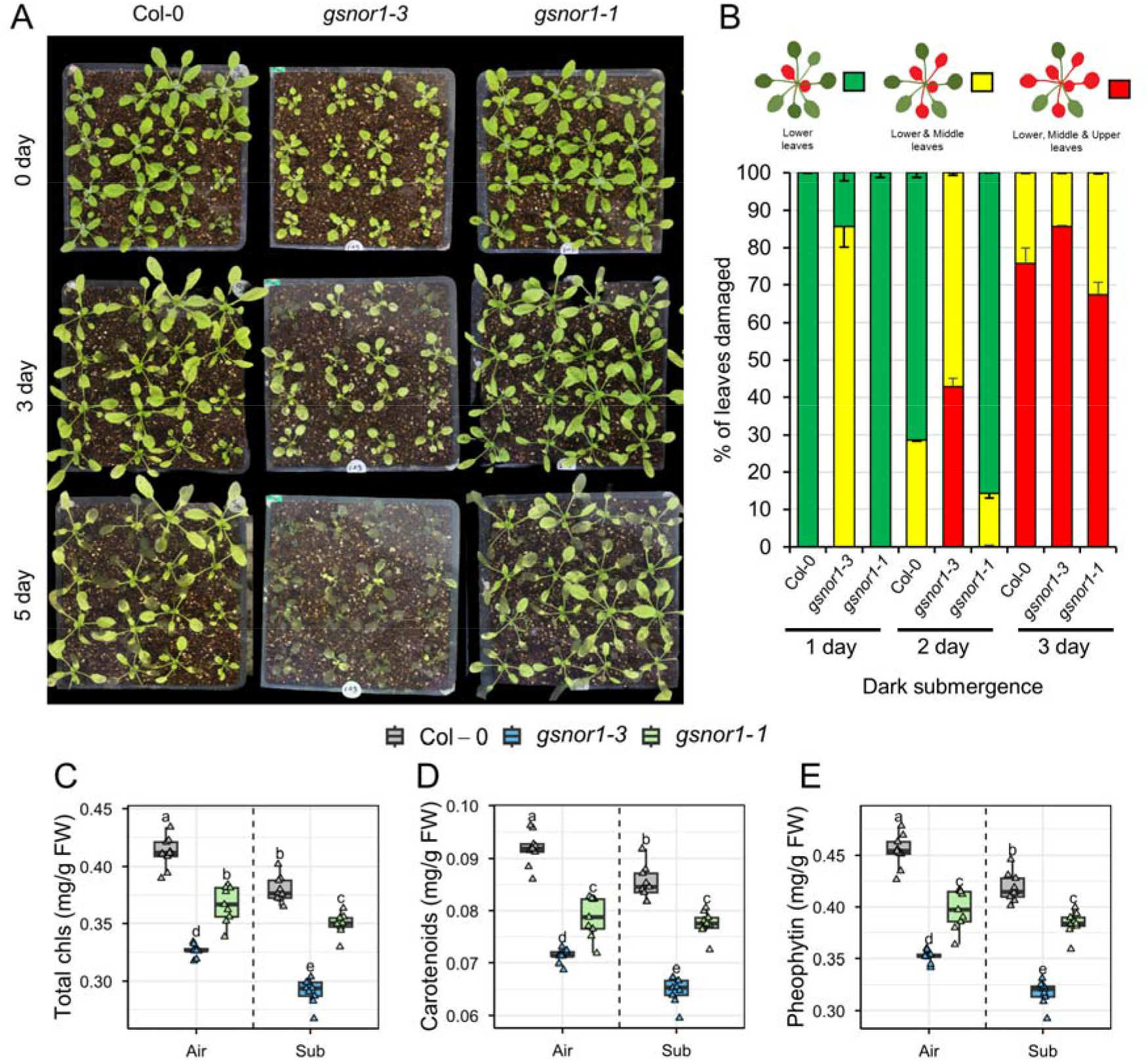
The role of S-nitrosoglutathione reductase 1 (GSNOR1) in the regulation of plant submergence tolerance. (A) Four-week-old Col-0, *gsnor1-3*, and *gsnor1-1* plants before submergence (day 0) and after 3- and 5-day dark submergence treatment. (B) Leaf damage assessment between Col-0, *gsnor1-3*, and *gsnor1-1* based on the number of decayed leaves per plant. Means (±SEs) are calculated from 5 individual plants. (C) Total chlorophyll (chls), (B) carotenoids, and (C) pheophytin levels in Col-0, *gsnor1-3*, and *gsnor1-1* plants exposed to air or 2 days of dark submergence (means ± SE, *n* = 9 technical replicates from 3 biological replicates). Different letters indicate statistically significant differences by two-way ANOVA and Tukey’s HSD (*P* < 0.05).

Additionally, we examined whether the intolerant phenotype of the *gsnor1-3* plant with elevated GSNO levels is linked to the N-degron pathway that may enhance proteolytic degradation. We collected leaf samples from Col-0, *gsnor1-3*, and *gsnor1-1* a day after submergence stress to measure transcriptional activation of enzymes involved in the N-degron pathway. qRT-PCR analysis showed lower gene expression levels of *PCO1*, *PCO2*, *ATE1*, and *PRT6* in both *gsnor1-3* and *gsnor1-1*, compared to Col-0, when plants grown in air (Figure 2A). Conversely, these genes were significantly upregulated in *gsnor1-3* under submergence, particularly *PCO2*, *ATE1*, and *PRT6*, compared to Col-0. In *gsnor1-1*, the expression of N-degron genes was either comparable to Col-0 or decreased under submergence stress (Figure 2A). To further assess the potential stabilization of ERFVII in the presence or absence of GSNO during dark submergence, we examined the gene expression of *RAP2.2*, *RAP2.3*, *RAP2.12*, and *HRE1*. Unlike the increased expression of enzymes involved in the N-degron pathway, ERFVIIs were downregulated in *gsnor1-3*, whereas ERFVIIs remained comparable in *gsnor1-1* and Col-0 (Figure 2B). Overall, these findings suggest that elevated GSNO/NO drives protein degradation via the N-degron pathway by altering cellular NO levels.

**Figure 2.**
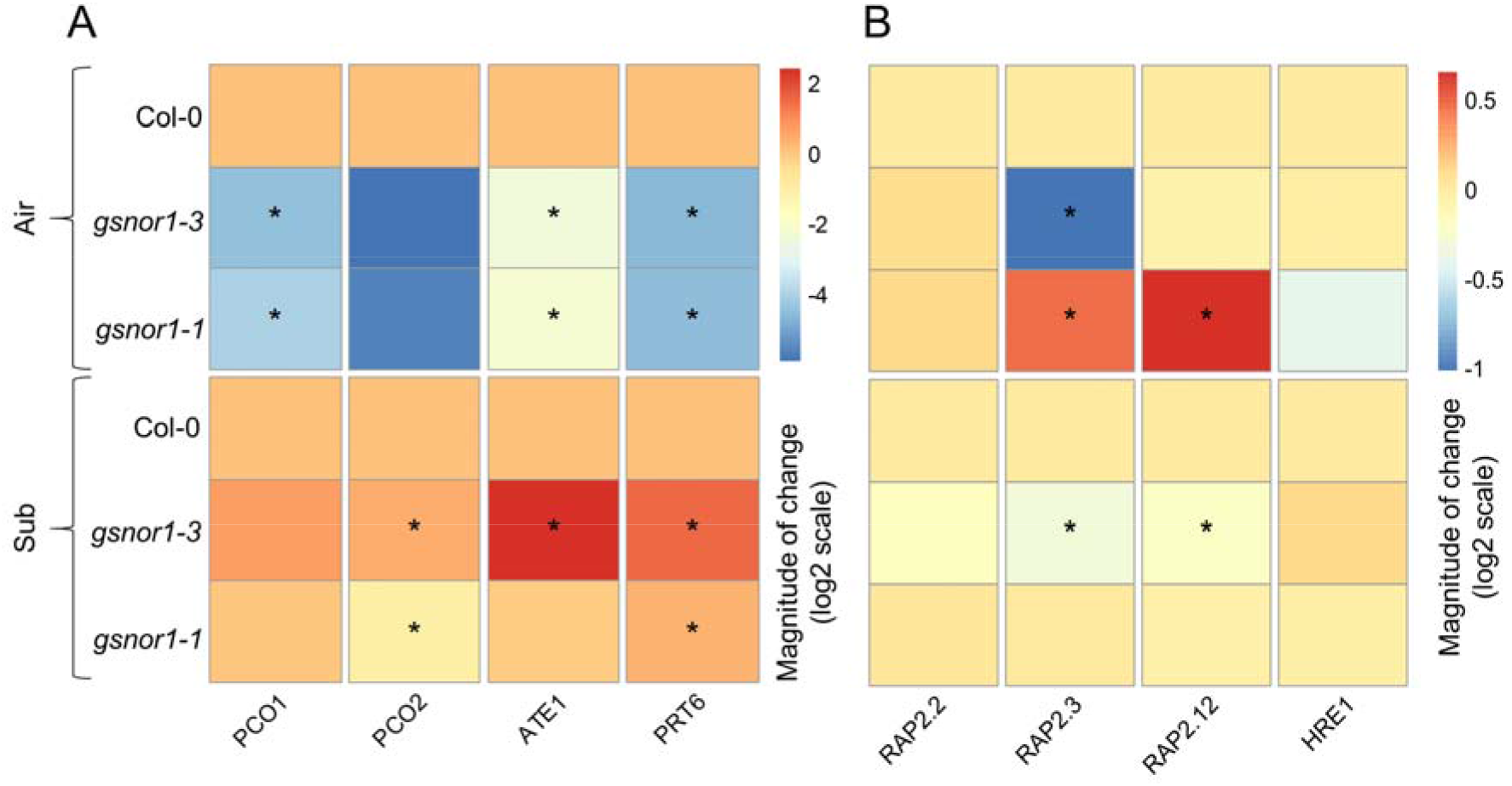
Expression pattern of genes involved in (A) the N-degron pathway and (B) ERFVII transcription factors. Heatmap representation of relative gene expression, assessed by qualitative real-time polymerase chain reaction (qRT-PCR), in Col-0, *gsnor1-3*, and *gsnor1-1* after 24 h of dark submergence. Means (±SEs) are calculated from 3 biological replicates, and the heatmap is displayed on a log2 scale. Asterisks indicate statistically significant differences by two-tailed Student *t*-test (\**p* < 0.05, \*\**p*< 0.01, \*\*\**p*< 0.01). ERFVII, group-VII ethylene response factor.

### The group VII ERF TFs undergo degradation via S-nitrosylation at Cys-2

As we know, group VII ERF TFs contain a conserved Nt-Cysteine (Cys) residue after methionine, which is oxidized in the presence of NO/O_2_ (Figure 3A). However, it remains unclear how NO-mediated non-enzymatic/enzymatic oxidation occurs. We hypothesized that S-nitrosylation may be involved in the oxidation process. To examine the potential Cys-mediated degradation of group VII via S-nitrosylation, we first used the GPS-SNO 1.0 tool to predict possible S-nitrosylation sites. GPS-SNO 1.0 prediction has an accuracy of 75.80%, with a sensitivity of 53.57% and a specificity of 80.14% ^29^. Intriguingly, we observed that five of the ERFVII have a high probability of being S-nitrosylated at Cys-2, as indicated by scores above 6 under the high threshold (Figure 3B). With this preliminary understanding, we aimed to validate a potential *in-silico* redox modification that could enhance ERFVII degradation. To confirm this, we modified the 2^nd^ Cys-SH to Cys-SNO in the five ERFVII using Schrödinger Suite 2024-4 (academic version) (Figure 4A-E). Changing the redox state of the group VII ERF TFs caused significant structural conformational changes, which are clearly visible in their overlapped structure of Cys-SH and Cys-SNO (Figure 4A-E). These changes suggest that NO signaling may largely influence the oxidation process.

**Figure 3.**
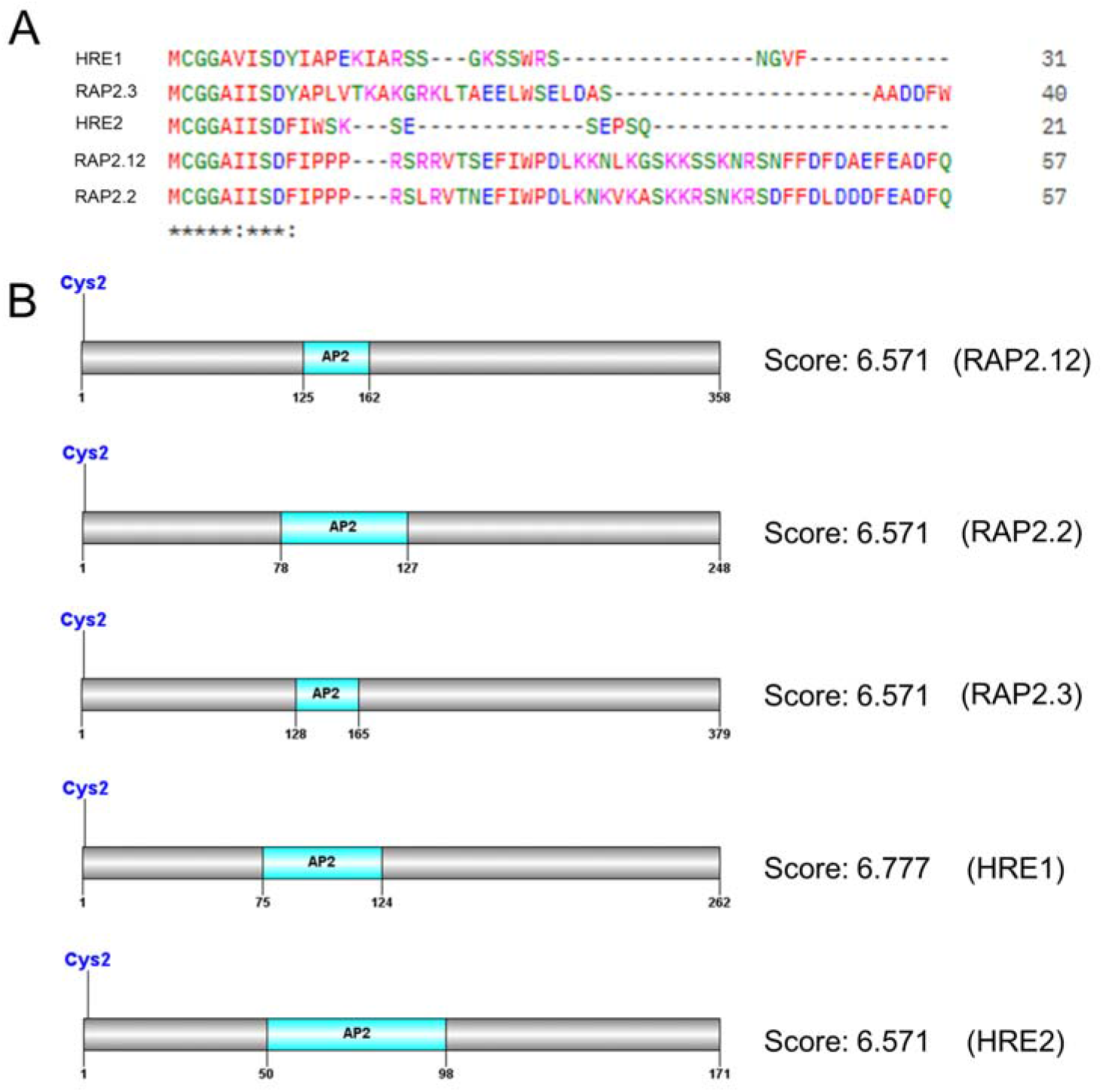
The group ERFVII TFs of *Arabidopsis thaliana*. (A) N-terminal sequence alignment of the group ERFVII TFs using CLUSTAL 0 (1.2.4) for multiple sequence alignment. (B) Predicted potential S-nitrosylation site at Cys-2 of the group ERFVII TFs using GPS-SNO 1.0. ERFVII, group-VII ethylene response factor; TF, transcription factor.

**Figure 4.**
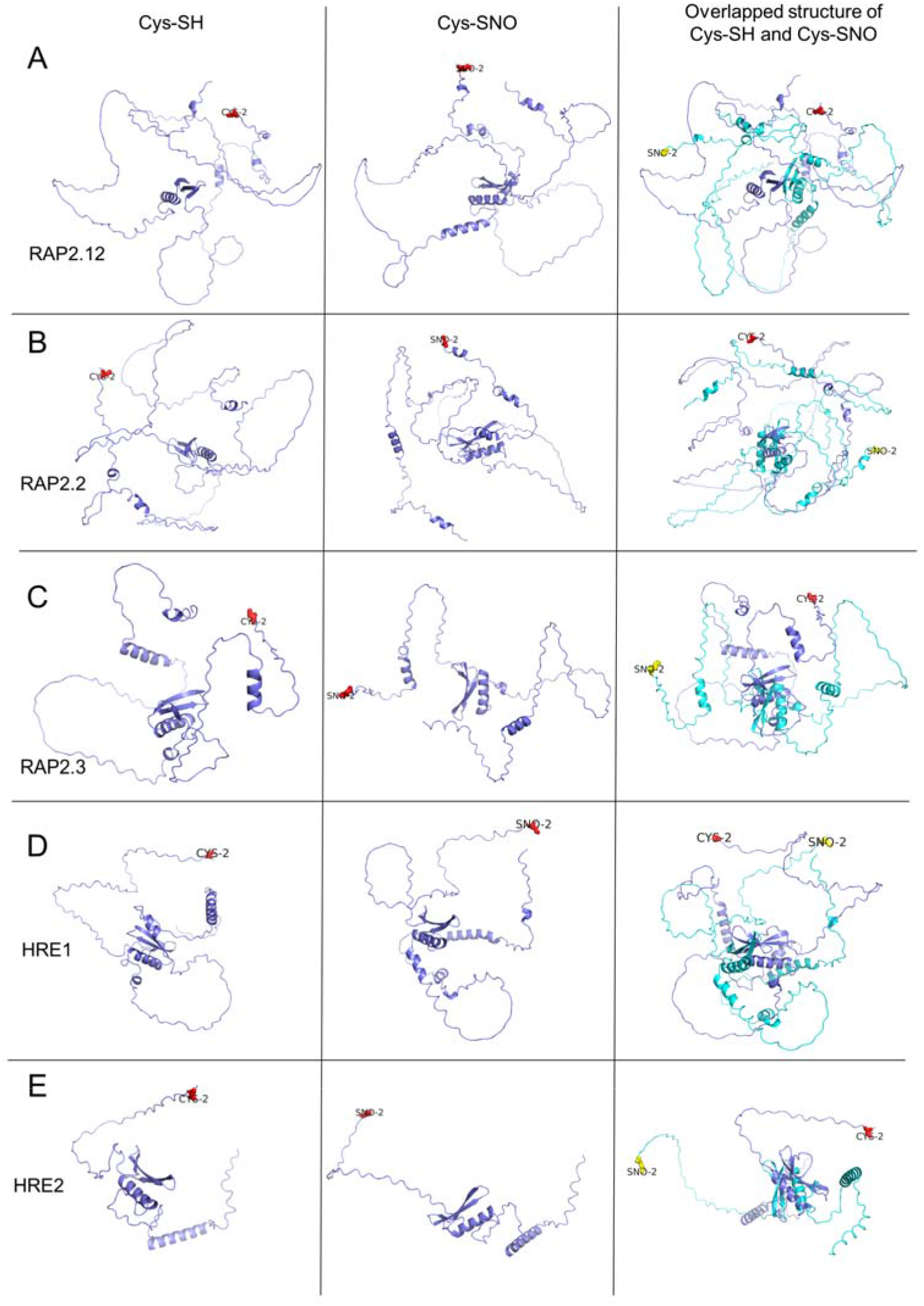
Alteration of the redox state in group ERFVII proteins, (A) RAP2.12, (B) RAP2.2, (C) RAP2.3, (D) HRE1, and (E) HRE2, facilitates the conformational changes. The second cysteine residue (Cys–SH) in each protein was structurally altered to its S-nitrosylated form (Cys–SNO) using Schrödinger Suite 2024-4 (academic version). The overlapping structures of Cys-SH and Cys-SNO are shown in purple and light blue, respectively. ERFVII, group-VII ethylene response factor; TF, transcription factor; RAP2.12/2.2/2.3, related to apetala 2.12/2.2/2.3; HRE1/2, hypoxia-responsive ERF.

Then, we assessed the interaction between the redox-modified ERFVII TFs and the enzyme that regulates the Arg/Cys branch of the N-degron pathway after oxidation. To do this, molecular docking of five group VII ERF isoforms (RAP2.12, RAP2.2, RAP2.3, HRE1, and HRE2) against the peptides of enzymes involved in the N-degron pathway (PCO1, PCO2, ATE1, and PRT6) was performed under both reduced (Cys-SH) and S-nitrosylated (Cys-SNO) redox states (Figure 5; Supplementary figure S2-4). Altered redox states of ERFVII isoforms-PCO1 docking was also performed, but no interaction was noticed at Cys2. For Peptides-PCO2, docking scores ranged from -3.982 to -5.903, with HRE2 exhibiting the highest binding affinity in both redox states (−5.903 and -5.034, respectively) (Figure 5A). In the Peptides-ATE1 dataset, overall docking scores were markedly improved across all isoforms (ranging from -6.61 to -7.090), while HRE2 again demonstrated the strongest binding affinity, particularly in the Cys-SNO state (−7.090) (Figure 5A). For Peptides-PRT6, docking scores ranged from -4.823 to -5.237, with comparable affinities observed across isoforms (Figure 5A). Notably, *in silico* redox-state alteration largely modulated docking scores across all three peptide-protein interactions, with HRE2-ATE1 consistently displaying the most favorable binding energetics, identifying it as the optimal interaction pair for further analysis. As PCOs are the first enzyme that precisely regulates the ERFVII degradation in the presence of O_2_, we were more curious about the changes of redox-modified ERFVII and PCO2.

**Figure 5.**
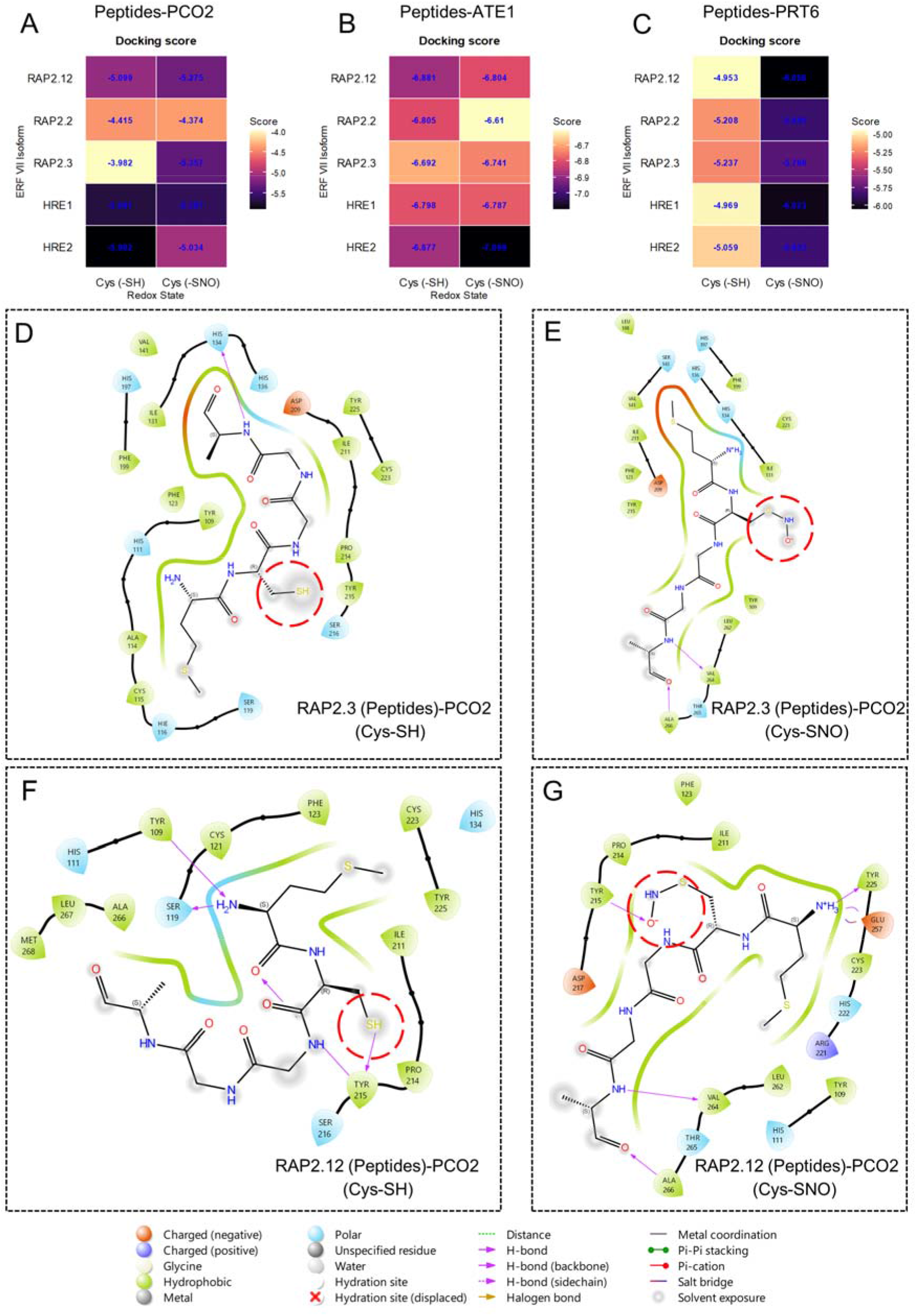
Interactions between peptides of group ERFVII (with or without redox modification) and N-degron pathway-related proteins. (A–C) Heatmap representation of docking scores between proteins and peptides under both redox states. With the best docked score, the 2D peptides-protein interaction of (D, E) RAP2.3 (Peptides)-PCO2 and (D, E) RAP2.12 (Peptides)-PCO2 are shown with or without redox modifications using Schrödinger Suite 2024-4 (academic version). ERFVII, group-VII ethylene response factor; RAP2.12/2.3, related to apetala 2.12/2.3; PCO2, plant cysteine oxidase.

Two-dimensional interaction diagrams of enzymes involved in the N-degron pathway with Peptides-ERFVII under Cys-SH and Cys-SNO states revealed distinct binding modes (Figure 5; Supplementary figure S2-4). The RAP2.3–PCO2 complex’s ligand (Cys-SH) binds via backbone H-bonding to HIS134, stabilized by polar residues (HIS136, HIS197, HIS111, SER119, SER216) and hydrophobic contacts (VAL141, ILE131, PHE199, PHE123, TYR109, PRO214, TYR215, ILE211, CYS223, TYR225) (Figure 5D). Upon oxidation to Cys-SNO, the interaction network shifts, targeting VAL264 and ALA266, while residues such as LEU188, SER143, LEU262, and THR265 reorganize (Figure 5E). Similarly, the RAP2.12– PCO2 complex shows extensive contacts in the reduced state, including hydrogen bonds, Pi-Pi stacking with TYR215, and interactions with CYS121, PHE123, CYS223, TYR225, ILE211, PRO214, LEU267, and MET268 (Figure 5F). In the oxidized state, interactions with TYR215 are lost, replaced by backbone bonds with VAL264 and ALA266, and shifted toward charged residues (Figure 5G). These shifts suggest that *in silico* S-nitrosylation of the conserved Cys induces conformational changes at the binding interface, remodeling ligand contacts, and potentially affecting peptide recognition in the presence of NO.

### Redox modification influences metabolic processes

Plants’ survival during submergence relies on their energy metabolism. We were curious how the GSNO-mediated redox status is involved in plant energy status under O_2_-deficient conditions. To determine the endogenous starch detection, levels of total soluble sugar, total carbohydrate levels, and relative gene expression of sugar-starvation markers *DARK INDUCIBLE 1* (*DIN1*) and *DIN6*. Results showed that *gsnor1-3* plants had reduced starch levels and reduced expression of *DIN1* and *DIN6* under dark submergence (Figure 6A, B). Moreover, *gsnor1-3* had low total soluble sugars and carbohydrate content under dark submergence, compared to Col-0 and gsnor1-1 (Figure 6C, D), suggesting that elevated GSNO levels compromised energy metabolism and dysregulated the canonical energy-starvation network, as evidenced by altered transcriptional activation of *DIN1* and *DIN6*, as well as of the energy-associated genes *MITOCHONDRIAL ATP SYNTHASE BETA-SUBUNIT* (*At5G08670*) and *HYPOXIA-RESPONSIVE MODULATOR 1* (*HRM1*) but no significant change in the expression levels of *ADP/ATP CARRIER 1* (*AAC1*) (Supplementary figure 5). Overall, these findings suggest that excessive GSNO levels failed to initiate a compensatory metabolic switch.

**Figure 6.**
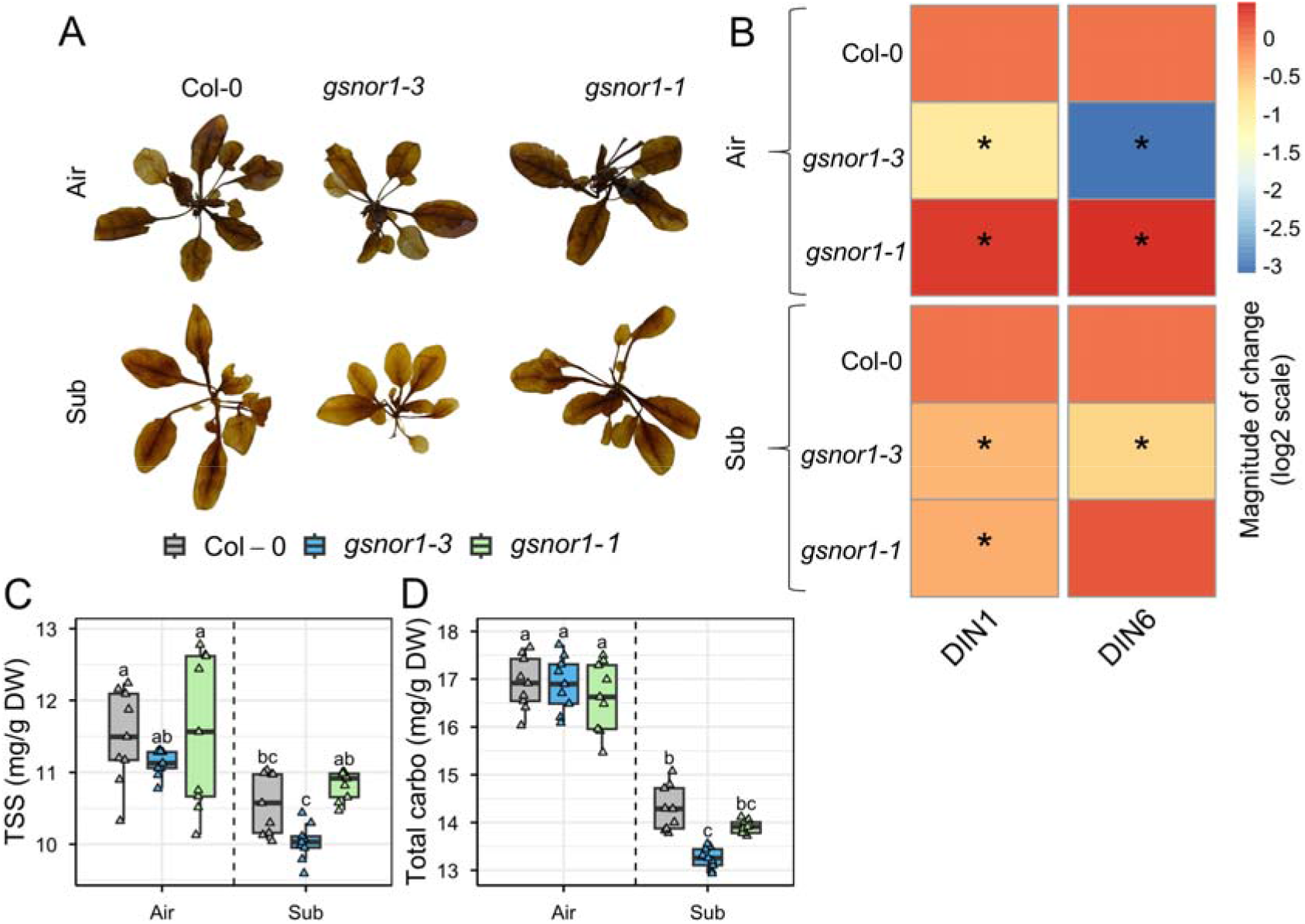
Impact of GSNO on energy metabolism under submergence. (A) Iodine-stained starch detection in leaves from Col-0, *gsnor1-3*, and *gsnor1-1* plants exposed to air or 24 h of dark submergence. (B) Heatmap representation of relative gene expression of carbon starvation marker genes *DIN1* and *DIN6*, assessed by qualitative real-time polymerase chain reaction (qRT-PCR), in Col-0, *gsnor1-3*, and *gsnor1-1* after 24 h of dark submergence (means ± SE, *n* = 3 biological replicates, and the heatmap is displayed on a log2 scale). (C) Total soluble sugar (TSS) and (D) total carbohydrate (carbo) levels in Col-0, *gsnor1-3*, and *gsnor1-1* plants exposed to air or 2 days of dark submergence (means ± SE, *n* = 9 technical replicates from 3 biological replicates). Asterisks indicate statistically significant differences by two-tailed Student *t*-test (\**p* < 0.05, \*\**p*< 0.01, \*\*\**p*< 0.01). Different letters indicate statistically significant differences by two-way ANOVA and Tukey’s HSD (*P* < 0.05). GSNO, S-nitrosoglutathione; DIN1/6, dark inducible 1/6.

With an understanding of NO responses during the energy crisis, we are interested in how redox modifications of group VII ERF TFs relate to their structural stabilization during metabolic reprogramming. To explore this, we assessed protein-ligand interactions between ERFVIIs and ATP. To do this, we performed molecular docking of five group VII ERF isoforms (RAP2.12, RAP2.2, RAP2.3, HRE1, and HRE2) against ATP under reduced (Cys-SH) and S-nitrosylated (Cys-SNO) redox states (Figure 7 and 8; Figure S6). Docking scores ranged from -2.148 to -4.135 across all isoforms and conditions. HRE2 exhibited the highest binding affinity in both redox states (Cys-SH: -4.135; Cys-SNO: -3.518), followed by HRE1 (Cys-SH: -3.383; Cys-SNO: -2.325) and RAP2.3 (Cys-SH: -3.937; Cys-SNO: -3.46). RAP2.12 and RAP2.2 displayed comparatively moderate affinities (Figure 7A). Overall, *in silico* redox modifications consistently reduced docking scores across most isoforms, suggesting that isoform-specific sensitivity to NO-mediated redox modifications with respect to ATP coordination.

**Figure 7.**
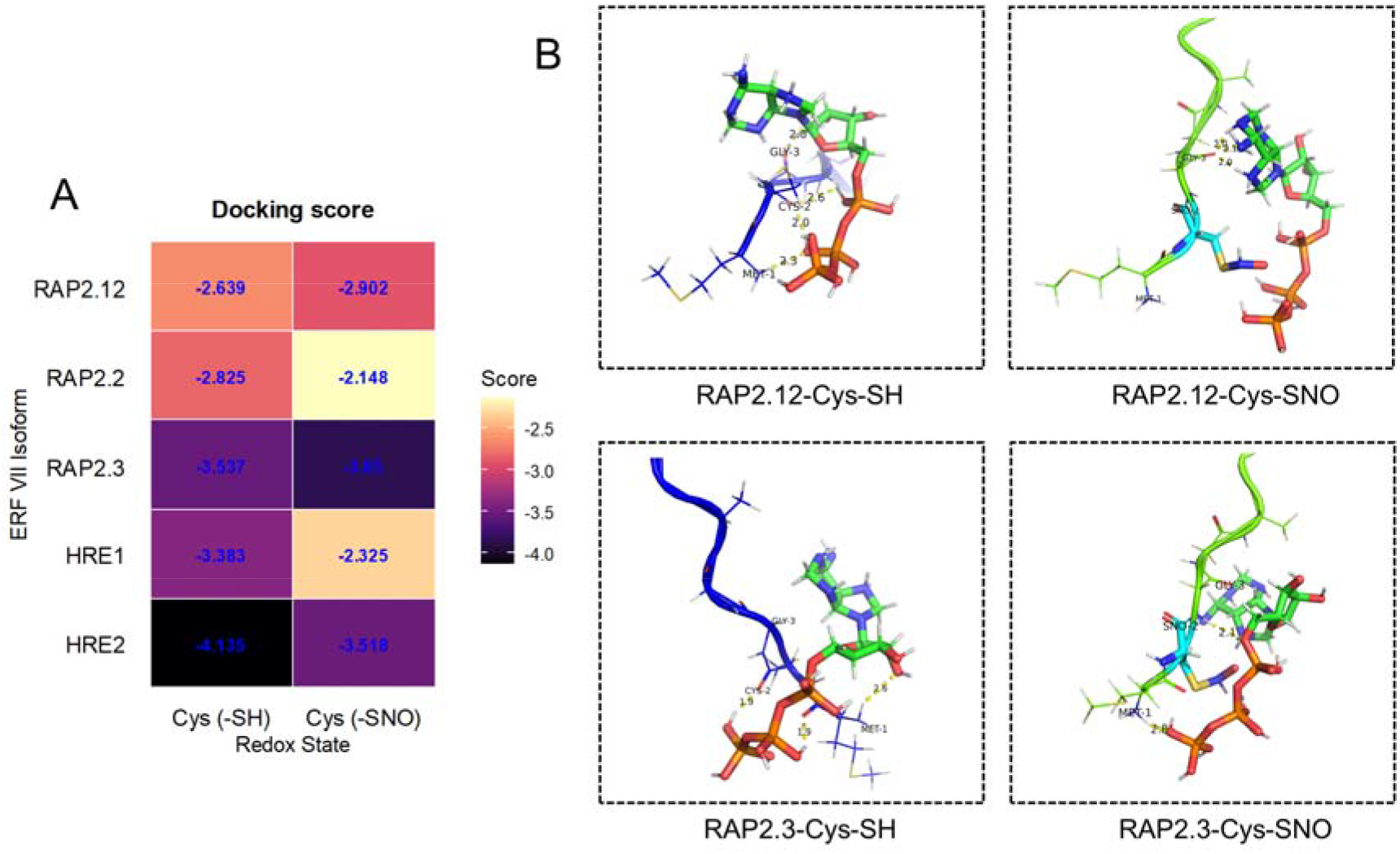
Protein-ligand interactions with or without redox modification. (A) Heatmap representation of docking scores between the group ERFVII proteins and ATP ligand under both redox states. (B) With the best docked score, the 3D protein-ligand interaction of RAP2.12-ATP and RAP2.3-ATP is shown with or without redox modifications using PyMOL 3.1.6.1 (academic version). ERFVII, group-VII ethylene response factor; RAP2.12/2.3, related to apetala 2.12/2.3.

Based on the favorable docking scores, 3D protein-ligand interactions of RAP2.12 and RAP2.3 with ATP were visualized under both redox states using PyMOL 3.1.6.1. In the reduced state (Cys-SH), both RAP2.12 and RAP2.3 displayed well-coordinated ATP binding, with key interactions involving conserved residues including CYS, GLY, and MET, at binding distances ranging from 1.9 to 2.6 Å, indicative of stable hydrogen bonding and electrostatic contacts (Figure 7B). The interaction networks were notably reorganized under oxidized state conditions (Cys-SNO), with altered binding geometries and reduced contact density, consistent with the attenuated docking scores observed in the heatmap. These structural observations collectively suggest that *in silico* S-nitrosylation of the conserved cysteine residue disrupts optimal ATP coordination, consistent with the attenuated docking scores, and suggest that the SNO modification repositions ATP into an alternative, potentially non-productive binding configuration.

To evaluate whether these docking-predicted differences reflect genuine conformational consequences of Cys-2 S-nitrosylation under dynamic conditions, we performed 100 ns molecular dynamics simulations of group VII ERFs under reduced (Cys-SH) and S-nitrosylated (Cys-SNO) redox states in the presence of ATP (Figure 8; Figure S7 and S8). Conformational sampling of RAP2.12-Cys-SH revealed a broadly distributed PCA landscape spanning PC1 values of approximately -75 to +125 Å and PC2 values of -75 to +50 Å, indicating extensive conformational flexibility. In contrast, RAP2.12-Cys-SNO displayed a more restricted sampling space concentrated along PC1 (−200 to +100 Å) with reduced PC2 dispersion (−75 to +50 Å), suggesting that the oxidized state constrains backbone mobility (Figure 8). RAP2.3-Cys-SH exhibited a tightly clustered distribution within PC1 (0 to +150 Å) and PC2 (−75 to +75 Å), reflecting conformational stability, whereas RAP2.3-Cys-SNO showed expanded sampling across PC1 (−60 to +30 Å) and PC2 (−150 to +75 Å), implying redox-induced structural perturbation (Figure 8).

**Figure 8.**
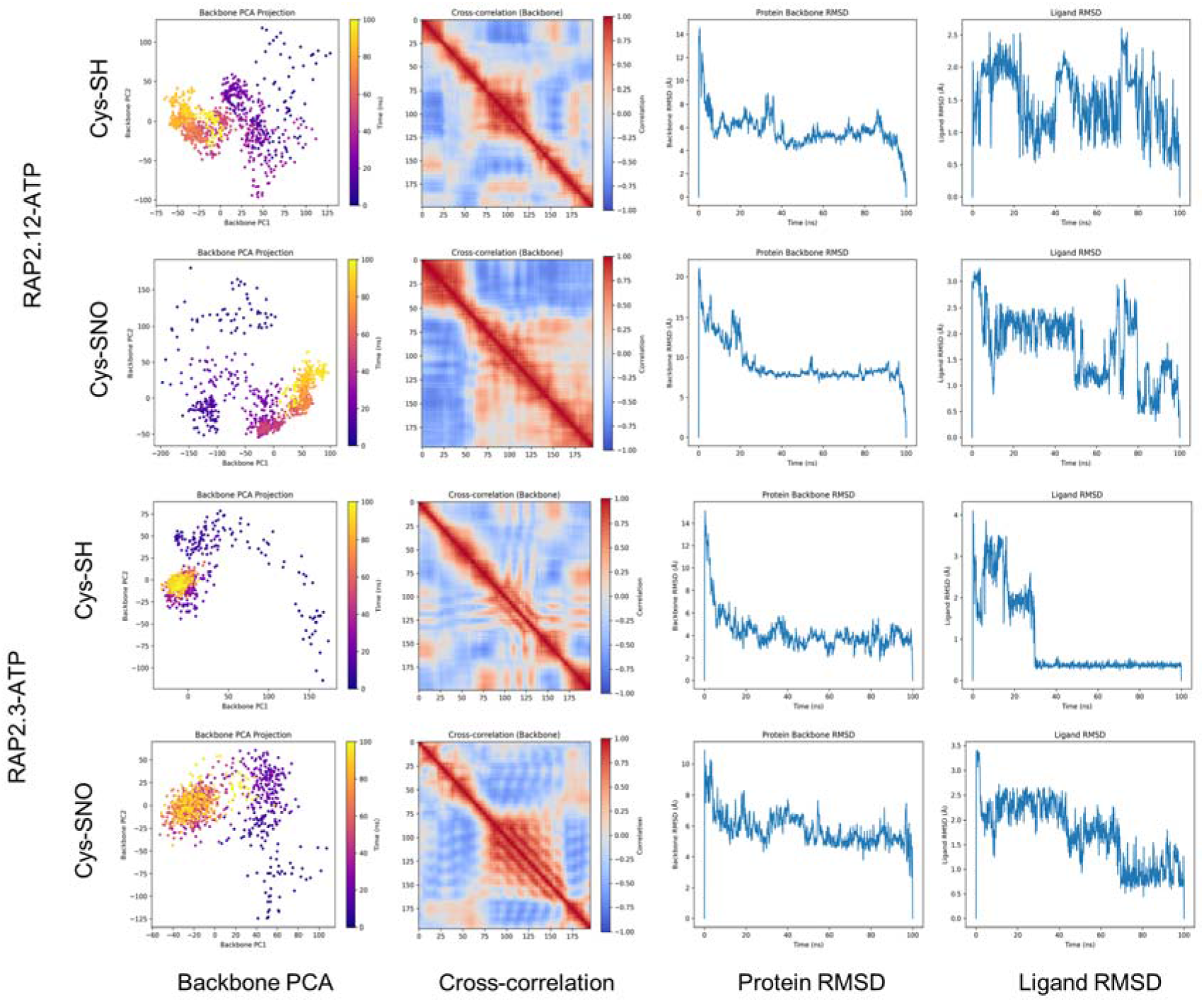
Molecular dynamics simulation analysis of RAP2.12-ATP and RAP2.3-ATP under reduced (Cys-SH) and S-nitrosylated (Cys-SNO) redox states. Backbone PCA projection showing conformational sampling, colored by time (purple = early, yellow = late). Dynamic cross-correlation matrices of backbone motions; red = correlated, blue = anti-correlated residue pairs. Protein backbone RMSD over time, reflecting structural stability relative to the initial docked conformation. Ligand (ATP) RMSD over time, indicating binding pocket stability throughout the simulation. ERFVII, group-VII ethylene response factor; RAP2.12/2.3, related to apetala 2.12/2.3; ATP, adenosine triphosphate; PCA, principal component analysis.

**Figure 9.**
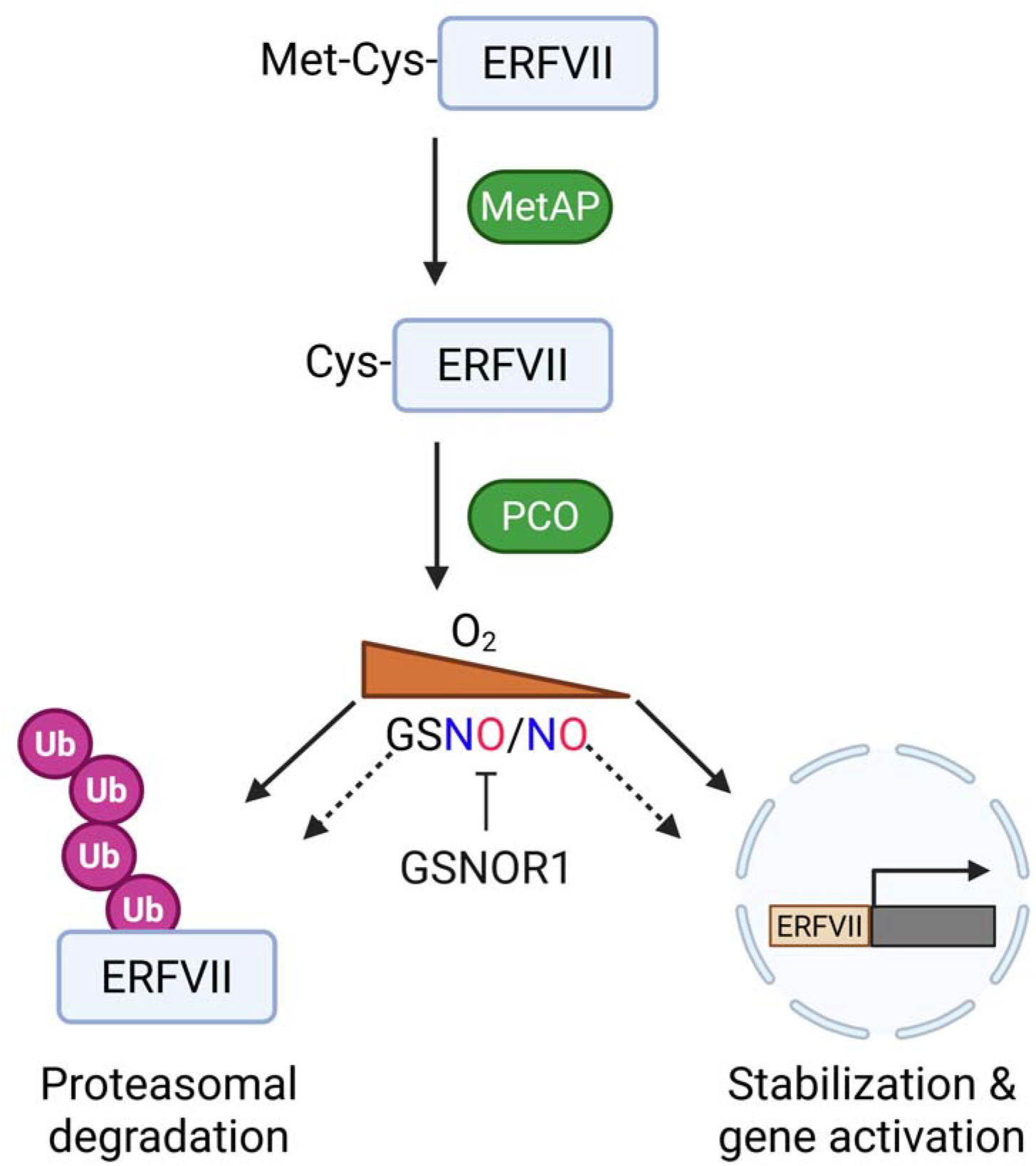
A putative model of the N-degron pathway that leads to ERFVII degradation via the 26S proteasome in the presence of O_2_/PCO/GSNO. Genetic and *in silico* analyses suggest that elevated GSNO/NO levels potentiate ERFVII degradation, while GSNOR1 controls endogenous GSNO levels, thereby altering degradation and stabilizing and activating hypoxia-marker genes. Met, methionine; Cys, cysteine; ERFVII, group VII ethylene response transcription factors; MetAP, methionine aminopeptidase; PCO, plant cysteine oxidase; GSNO, S-nitrosoglutathione; GSNOR1, S-nitrosoglutathione reductase 1.

The cross-correlation matrices of RAP2.12-Cys-SH displayed predominantly correlated backbone motions (correlation values approaching +1.00) concentrated along the diagonal, while RAP2.12-Cys-SNO exhibited increased anti-correlated regions (values reaching -1.00), indicative of disrupted intramolecular communication. RAP2.3-Cys-SH showed a fragmented correlation pattern with pronounced anti-correlated domains, which became more structured and regionally correlated upon S-nitrosylation, suggesting redox-dependent reorganization of residue-residue coupling networks across the ∼180-residue backbone (Figure 8). Moreover, RAP2.12-Cys-SH exhibited initial structural fluctuations ranging from ∼4 to 14 Å during the first 40 ns, progressively stabilizing to ∼4–6 Å toward the end of the simulation. The Cys-SNO state maintained moderate fluctuations between ∼4 and 10 Å throughout the 100 ns trajectory. RAP2.3-Cys-SH demonstrated higher initial RMSD values of ∼10–14 Å that gradually equilibrated to ∼4–6 Å after 60 ns, whereas RAP2.3-Cys-SNO stabilized at comparatively lower RMSD values of ∼2–6 Å, suggesting that S-nitrosylation promotes structural compaction in RAP2.3 (Figure 8).

ATP binding stability differed markedly between redox states. In RAP2.12-Cys-SH, ligand RMSD fluctuated considerably between ∼1.0 and 2.5 Å throughout the simulation, whereas the Cys-SNO state displayed an initial displacement of ∼2.5–3.0 Å that gradually declined to ∼1.0–1.5 Å toward 100 ns, indicating progressive ligand stabilization. In RAP2.3-Cys-SH, ATP exhibited high initial displacement of ∼2.5–4.0 Å during the first 20 ns, followed by convergence to ∼0.5 Å, indicating stable binding after equilibration. The Cys-SNO state of RAP2.3 maintained a relatively stable ligand RMSD of ∼1.5–2.5 Å throughout, with minor fluctuations suggesting consistent ATP positioning within the binding pocket (Figure 8). Collectively, the docking, interaction visualization, and MD simulation data demonstrate that Cys2 S-nitrosylation across ERFVII isoforms generally attenuates predicted ATP binding affinity while inducing isoform-specific conformational consequences, providing a structural basis for how NO-mediated redox modification of ERFVIIs may relate to the differential energy metabolism gene expression observed in *gsnor1-3* and *gsnor1-1* plants under submergence stress.

## Discussion

Ethylene-mediated NO depletion by PHYTOGLOBIN1 (PGB1) enhances the stability of ERFVIIs, thereby enabling plants to pre-adapt and survive under hypoxic conditions ^19^. Likewise, the extensive overlap in global gene expression profiles among NO-deficient mutants, N-degron pathway mutants, and hypoxia-treated wild-type plants supports a central role for NO in regulating ERFVII stability ^17^. Although previous studies established that both O_2_ and NO are required for ERFVII degradation, the molecular sequence linking these signals remains unresolved. One long-standing hypothesis proposes that the conserved N-terminal Cys is first modified by NO and subsequently oxidized in an O_2_-dependent manner ^16^, yet direct experimental evidence for this mechanism has been lacking.

S-nitrosylation is an evolutionarily conserved NO-dependent post-translational modification that regulates protein functions by converting Cys thiols into SNO ^30^. Cellular S-nitrosylation is precisely controlled by GSNOR through the regulation of NO reservoir GSNO, and numerous studies have demonstrated its functional roles in plant development and stress tolerance ^24,31^. However, identifying S-nitrosylated proteins *in vivo* remains technically challenging because of their low abundance and intrinsic instability, despite advances in approaches such as the biotin-switch technique ^32^. In this study, we hypothesized that Nt-Cys oxidation of the Arabidopsis group VII ERF TFs, including RAP2.12, RAP2.2, RAP2.3, HRE1, and HRE2, may occur via S-nitrosylation. We used genetic analyses with structural modeling, molecular docking, and molecular dynamics simulations to test whether the conserved N-terminal Cys2 residue of Arabidopsis ERFVII proteins represents a potential target for NO-mediated S-nitrosylation.

Genetic evidence supported a negative relationship between elevated GSNO levels and submergence tolerance. The *gsnor1-3* mutant, which accumulates GSNO because of impaired GSNOR activity, exhibited pronounced sensitivity to prolonged dark submergence, whereas the gain-of-function *gsnor1-1* mutant displayed slightly improved tolerance relative to wild type (Figure 1). Elevated GSNO in *gsnor1-3* was accompanied by increased expression of N-degron pathway genes (*PCO1*, *PCO2*, *ATE1*, and *PRT6*) together with reduced transcript abundance of multiple ERFVII genes (Figure 2). These observations are consistent with enhanced NO-dependent turnover of ERFVII proteins and suggest that GSNO homeostasis influences both O_2_ sensing and hypoxia-responsive transcription.

Computational analyses further supported this model. All five Arabidopsis ERFVII proteins were predicted to contain a high-confidence S-nitrosylation site at the conserved N-terminal Cys2 residue. Although these predictions require biochemical validation, they are supported by the performance of GPS-SNO 1.0 and by previous experimental studies that validate its predictive capability ^29,33–35^. Structural modeling revealed that conversion of Cys-SH to Cys-SNO induced conformational rearrangements in every ERFVII protein, indicating that S-nitrosylation could alter protein structure before proteolytic degradation.

These altered redox states of five ERFVII were docked against the N-degron pathway genes (PCO2, ATE1, and PRT6). Among the ERFVII family members, HRE2 exhibited the strongest predicted response to S-nitrosylation (Figure 3; Figure 4; Figure S2-4). The remarkable improvement of ATE1 affinity (−7.090) under Cys-SNO and the recruitment of previously non-contacting residues (MET407, VAL448) upon oxidation indicate a very significant allosteric effect that results from the addition of the –SNO group. Under hypoxic conditions, HRE2 is reported to be regulated post-transcriptionally through mRNA stabilization and to be involved in anaerobic signaling, functioning as an efficient substrate for the N-degron pathway ^11,36^. The protein itself has been directly confirmed as a PRT6/BIG substrate, and it is also the one found in increased quantities in *prt6* big double mutants, with no change in transcripts ^37^. In *gsnor1-3*, chronic GSNO effects would be predicted to continuously nitrosylate newly synthesized HRE2, forming a loop that creates new proteins while targeting them for N-degron-mediated degradation. Thus, it leads to the depletion of an active pool of HRE2, which is normally employed in transcription precisely at the time when the limitation of O_2_ most needs its presence. Such a mechanism provides a plausible explanation for the reduced flooding tolerance observed in *gsnor1-3* compared with wild-type plants, which maintain a more balanced NO homeostasis during submergence ^38^.

Beyond regulating protein turnover, our data suggest that NO-mediated Cys2 S-nitrosylation may also influence energy metabolism under O_2_-deficient conditions. The hypersensitive phenotype of *gsnor1-3* may be due to a redox burst together with inefficient energy use during complete submergence. We found that *gsnor1-3*, with excess GSNO levels, compromised energy status during dark submergence, while their starvation marker genes were unlikely to be dysfunctional (Figure 6). On the other hand, increased energy demand in *gsnor1-3* under O_2_-deficient conditions upregulates transcript levels related to energy metabolism, but not in the *gsnor1-1* mutant (Supplementary figure S5). Upregulation of sugar-utilizing enzymes under low-O_2_ conditions is an energy-consuming process that balances between energy consumption and production ^39^. This genetic contrast prompted us to examine where the ERFVII redox state itself regulates the metabolic processes. Across all five ERFVII transcription factors, *in silico* S-nitrosylation at Cys2 consistently reduced ATP-binding affinity, with reorganization of the interaction network and reduced contact density observed in the Cys-SNO states. While ERFVII TFs are not canonical ATPases or nucleotide-binding enzymes, the interaction of AP2/ERF domain proteins with ATP and other nucleotides through a positively charged surface region is not without precedent, and such interactions have been proposed to modulate DNA-binding affinity and transcriptional activity ^40^. We therefore propose that the reduced ATP coordination capacity of Cys-SNO ERFVIIs impairs the recruitment of ATP-dependent chromatin remodeling complexes to hypoxia-responsive promoter elements, thereby suppressing anaerobic transcriptional programs ^41^. Under this mechanism, the elevated GSNO in *gsnor1-3* would continuously shift ERFVIIs toward the Cys-SNO state, impairing HRPE-driven gene activation and compelling compensatory transcriptional upregulation of mitochondrial energy metabolism genes (Figure 7).

Molecular dynamics simulations conducted over 100 ns further revealed that this modification induces conformational changes in the protein, affecting both its structural stability and the stability of ATP within the binding pocket (Figure 8; Supplementary figure 7 and 8). The molecular dynamics studies also corroborated the docking results: RAP2.12 with a free sulfhydryl (SH) on its cysteine moved easily into many shapes (covering -75 to +125 Å on PC1), and its backbone’s position became more and more stable over time. However, RAP2.12 with a nitrosylated Cys (SNO) was much more limited in the shapes it could adopt. RAP2.3 behaved in the opposite manner: the sulfhydryl form was tightly grouped and quickly reached balance, whereas the nitrosylated form showed a much wider spread of possible shapes. These isoform-specific responses parallel the differential effects of S-nitrosylation on K-RAS and hemoglobin, where attachment of NO to a single Cys produced opposite effects on backbone flexibility depending on the structural context of the modified site ^42^. Taken together, the structural data suggest that *in silico* Cys2 S-nitrosylation in ERFVIIs produces a dual molecular consequence: impaired ATP binding, which may reduce transcriptional competence prior to proteolysis, and a conformational bias toward states that are recognized more efficiently by N-degron pathway enzymes. Such a two-step mechanism would allow NO to fine-tune ERFVII activity by attenuating transcriptional function before proteasomal degradation.

The phenotypic and transcriptional behavior of *gsnor1-1* mutant provides complementary support for the *in silico* predictions. Enhanced GSNOR activity reduces GSNO accumulation and limits cellular S-nitrosylation, thereby maintaining lower expression of N-degron pathway genes, preserving hypoxia-responsive transcription, and improving submergence tolerance (Figure 7). If Cys2 S-nitrosylation is the initiating step in ERFVII proteolysis, lower SNO availability in *gsnor1-1* would be expected to preserve a greater fraction of ERFVII in the reduced, transcriptionally active Cys-SH conformation, exactly as the molecular dynamics data predict for the energetically accessible, broadly sampled conformational states characteristic of Cys-SH isoforms (Figure 4A-E). The broader PCA landscape of RAP2.12-Cys-SH and the rapid RMSD equilibration of RAP2.3-Cys-SH collectively point to a dynamically flexible protein poised for transcriptional engagement, in contrast to the more constrained Cys-SNO forms that preferentially populate binding-competent conformations for N-degron enzyme recognition. This correspondence between computed structural behavior and measured phenotypic outcomes lends credibility to the proposed mechanism and supports the use of *in silico* redox modification as a hypothesis-generating tool in situations where possible biochemical quantification of labile SNO intermediates remains technically challenging.

### Conclusions & limitations of the study

Our genetic and computational analyses support a model in which GSNOR1-mediated regulation of GSNO levels acts as a molecular rheostat controlling ERFVII stability and hypoxia-responsive signaling under O_2_ deficiency. Elevated NO/GSNO levels are predicted to promote S-nitrosylation of the conserved N-terminal Cys2 residue, inducing conformational changes that enhance interactions with components of the Arg/Cys branch of the N-degron pathway while perturbing ATP coordination and cellular energy homeostasis. These combined effects are associated with faster ERFVII degradation, reduced ERFVII gene expression, and a more severe energy crisis under flooding stress. Conversely, the lower levels of GSNO in *gsnor1-1* mutant plants are consistent with preservation of the reduced Cys-SH state, stabilization of ERFVII proteins, sustained anaerobic gene expression, and enhanced adaptation to O_2_ deficiency.

Our findings extend the current model that both O_2_ and NO are required for ERFVII degradation ^43^, by proposing that S-nitrosylation of the conserved N-terminal cysteine may represent an early NO-dependent redox event that facilitates subsequent PCO-mediated Cys oxidation and N-degron recognition. However, this mechanistic model remains provisional because the *in silico* Cys-SNO substitution represents a structural approximation and cannot fully capture the complexity of the cellular redox environment, including competing cysteine oxidation states, glutaredoxin- and thioredoxin-mediated redox buffering, or the spatial dynamics of NO production ^44^. Future biochemical validation, including biotin-switch assays to detect ERFVII S-nitrosylation, site-directed mutagenesis of the conserved Cys2 residue, and co-immunoprecipitation analyses of ERFVII interactions with PCO2 and ATE1 under defined NO conditions, will be essential to test the proposed mechanism. Together, this work provides a framework for understanding how NO-mediated redox signaling integrates with O_2_ sensing to regulate ERFVII stability and flooding adaptation in plants.

## METHOD DETAILS

### Plant materials and experimental design

All mutants and transgenic lines for this study were sourced from the *Arabidopsis thaliana* Columbia (Col-0) ecotype. The T-DNA mutants *gsnor1-1* and *gsnor1-3*, with gain- and loss-of-function mutations in GSNOR1, respectively, were described previously ^22^. Flooding tolerance assays were performed as described by Licausi et al. ^45^. Briefly, four-week-old plants were submerged in 23-cm-high plastic boxes and kept in the dark. Rosettes were submerged to a depth of 14 cm below the water surface. The photograph was taken above the water level on the indicated days. To quantify the damage, wilting and necrotic leaves were counted following the methodology of Tsai et al. ^46^.

### Assessment of photosynthetic pigments

Total chlorophyll, carotenoids, and pheophytin were quantified from 80% acetone extracts by measuring absorbance at 663, 645, and 470 nm using a UV spectrophotometer (Thermo Scientific™ Multiskan™ GO Microplate Spectrophotometer, Ratastie, Finland). Chlorophylls, carotenoids, and pheophytin were calculated according to Arnon ^47^, Lichtenthaler and Wellburn ^48^, and Lichtenthaler ^49^, respectively, based on sample weight and acetone volume.

### Quantification of H_2_O_2_ and GSH levels

H_2_O_2_ levels in Arabidopsis rosette leaves were measured as described by Tomassi et al. ^50^. The level of reduced glutathione (GSH) was estimated according to Ellman ^51^ with modifications. Freeze-dried samples (5 mg) were extracted with 0.8 mL 5% (v/v) trichloroacetic acid (TCA) and centrifuged at 14,000 rpm for 20 min at 4 °C. 100 μL supernatant was added with 1 mL 150 mM potassium-phosphate (KP) buffer (pH 7) and 150 μL 5,5’-dithiobis (2-nitrobenzoic acid) (DTNB) (∼ 20 mM, 75.3 mg in 30 mL 100 mM KP buffer; pH 7). This mixture was then incubated at 30°C for 5-10 minutes, and absorbance was measured at 412 nm. GSH content was calculated using a standard curve of 1 mg/100 mL GSH ranging from 0 to 10 µM and expressed as µM/g DW.

### RNA isolation and qRT-PCR analysis

Total RNA was extracted from 200 mg of liquid nitrogen-ground samples. Homogenization was performed with 1 mL of Trizol reagent, followed by the addition of chloroform and isopropanol, and the mixture was washed with 500 μL of 70% ethanol. cDNA synthesis and qRT-PCR were performed following the protocol of Das et al. ^52^. *UBQ5* served as the reference gene. Primers are listed in Supplementary Table 1.

### Iodine staining and measurement of total soluble sugars and total carbohydrates

For iodine staining, Arabidopsis plants were collected and immersed in 95% ethanol to extract chlorophyll. After washing with deionized water, the leaves were stained with Lugol’s iodine solution for 10□min, and representative plants from three biological replicates were photographed accordingly ^53^. Total soluble sugar (TSS) and total carbohydrate content were measured using the core methodologies of Somogyi ^54^ and DuBois et al. ^55^, respectively, with slight modifications as described by Das et al. ^52^.

### Multiple sequence alignment and prediction of S-nitrosylation site

Protein sequences were aligned using the European Molecular Biology Laboratory-European Bioinformatics Institute Clustal Omega MSA tool (CLUSTAL 0(1.2.4)) ^56^. We used GPS-SNO 1.0 software to predict and identify cysteine residues that could be targets of NO-based S-Nitrosylation ^29^. GPS-SNO 1.0 has demonstrated efficiency and speed in predicting S-Nitrosylation sites, achieving an accuracy of 75.80%, a specificity of 53.57%, and a sensitivity of 80.14%.

### Structure prediction using AlphaFold 3

The amino acid sequences of target proteins, RAP2.12 (AT1G53910), RAP2.3 (AT3G16770), RAP2.2 (AT3G14230), HRE1 (AT1G72360), HRE2 (AT2G47520), PCO1 (AT5G15120), PCO2 (AT5G39890), ATE1 (AT5G05700), and PRT6 (AT5G02310), were retrieved from the Tair database (https://www.arabidopsis.org/) using their respective accession numbers. The three-dimensional structures of proteins were predicted using AlphaFold 3, which employs deep learning models to accurately predict protein and protein–ligand complex structures ^57,58^. Model quality was assessed using pLDDT confidence metrics, and the most reliable structures were selected for downstream analyses.

### Structural modification for S-nitrosylation

To simulate post-translational modification, the second cysteine residue (C–SH) in each protein was structurally altered to its S-nitrosylated form (C–SNO) using Schrödinger Suite 2024-4 (academic version) builder option. Both unmodified and modified sequences were then resubmitted to AlphaFold 3 to predict their modified conformations ^57,58^.

### Small-molecule docking

Docking of small-molecule ligands was performed using AutoDock Vina (v1.1.2) to predict their binding affinity and orientation toward the target protein ^59^. A cubic grid box was centered on the protein’s active site, covering all residues within the defined binding region to allow unrestricted ligand movement. The exhaustiveness parameter was set to 8 to ensure adequate conformational sampling. The docking runs were performed using the Lamarckian genetic algorithm, and the pose with the lowest binding energy (kcal/mol) was selected as the most favorable conformation. The resulting protein–ligand complex was visually examined and exported as a PDB file for subsequent molecular dynamics simulations.

### Peptide docking

For the peptide docking experiments, AutoDock CrankPep (ADCP v1.1) was employed to accommodate the intrinsic flexibility of peptide ligands ^60^. This algorithm integrates fragment-based assembly with conformational sampling to accurately predict peptide– receptor interactions. The receptor was treated as rigid, while full flexibility was allowed for the peptide backbone and side chains. The docking results were ranked by their binding energy scores, and the top-scoring peptide conformation was selected for further refinement. All docking outputs, including both small-molecule and peptide complexes, were visualized and analyzed using Schrödinger Maestro (academic version, 2024-4 release) and Discovery Studio Visualizer (academic version, 2025 release) to inspect hydrogen bonds, hydrophobic interactions, and surface complementarity within the binding site.

### Protein and ligand preparation

ATP ligand was obtained from PubChem (CID: 5957). The receptor and ligand structures were prepared using AutoDockTools (ADT) prior to molecular docking. All water molecules, cofactors, and crystallographic heteroatoms not involved in the binding process were removed. Polar hydrogens were added, and Gasteiger charges were assigned to ensure proper electrostatic representation of the system. The prepared receptor structure was saved in PDBQT format, while ligand molecules were geometry-optimized using internal energy minimization algorithms to eliminate steric clashes and obtain low-energy conformers suitable for docking.

### Molecular dynamics simulation

#### System preparation

The selected docked complexes were subjected to molecular dynamics (MD) simulations using Desmond v8.0 ^61^, part of the Schrödinger 2024-4 Academic Suite. Each complex was solvated in an orthorhombic box with TIP3P water molecules, ensuring a 10 Å buffer distance between the solute and box boundaries. To maintain charge neutrality, appropriate counterions (Na□ and Cl□) were added, and an additional 0.15 M NaCl solution was introduced to emulate physiological conditions. The systems were parameterized using the OPLS_2005 force field for both the protein and ligand components.

#### Energy minimization and equilibration

Energy minimization was performed using the steepest descent algorithm for 2000 iterations or until the system’s energy converged. Following minimization, the system underwent a two-phase equilibration: first under the NVT ensemble (constant particle number, volume, and temperature) at 300 K for 1 ns using the Berendsen thermostat, and subsequently under the NPT ensemble (constant pressure and temperature) at 1 atm for 1 ns using the Berendsen barostat. These stages ensured temperature and pressure stabilization before commencing the production run.

#### Production dynamics

The production MD simulations were conducted for 100 ns with a 2-fs integration time step, applying periodic boundary conditions in all dimensions. Long-range electrostatic interactions were treated using the Particle Mesh Ewald (PME) method, while the SHAKE algorithm was used to constrain all covalent bonds involving hydrogen atoms. Trajectories were saved every 100 ps for subsequent analyses, producing approximately 1000 frames per simulation. Throughout the simulation, the temperature was maintained at 300 K and the pressure at 1 atm, representing near-physiological conditions.

### Post-simulation analysis

#### Trajectory processing

The trajectory and topology files obtained from Desmond were analyzed using Python 3.10 with scientific and bioinformatics libraries, including MDAnalysis (v2.9.0), MDTraj (v1.10.3), Matplotlib (v3.10.7), Seaborn (v0.13.2), NumPy (v2.2.6), Pandas (v2.3.3), ProLIF (v2.0.3), and BioPython (v1.85). The analyses were performed in a Linux-based environment to ensure compatibility with Desmond-generated trajectory formats. Custom Python scripts automated data extraction, processing, and visualization to evaluate key dynamic and structural parameters across the entire trajectory.

#### Structural and dynamic analyses

The root-mean-square deviation (RMSD) of Cα atoms was computed to monitor the temporal stability of the protein backbone, while the radius of gyration (Rg) was used to assess the compactness of the overall structure. The root-mean-square fluctuation (RMSF) provided residue-level insights into local flexibility, allowing the identification of highly mobile or stabilized regions. Conformational sampling across the simulation was examined by generating pairwise RMSD matrices and performing principal component analysis (PCA) to identify dominant collective motions. Additionally, the dynamic cross-correlation matrix (DCCM) was computed to assess correlated and anticorrelated movements among residues, providing insights into coordinated domain motions.

#### Ligand and interaction stability

The ligand RMSD, RMSF, and radius of gyration were calculated to track conformational changes of the bound molecules throughout the simulation. The solvent accessible surface area (SASA) was evaluated using FreeSASA to assess fluctuations in solvent exposure over time. Hydrogen bond analysis was performed using MDAnalysis.analysis.hydrogenbonds.HydrogenBondAnalysis, applying a donor–acceptor distance cutoff of 3.5 Å and an angle cutoff of 120°. Hydrogen bond frequency and persistence were quantified, and residues forming persistent contacts were identified. In addition, a contact frequency analysis was performed to identify hydrophobic or aromatic residues within 4.0 Å of the ligand atoms, highlighting residues critical to maintaining binding stability.

#### Binding free energy calculation

To estimate the thermodynamic strength of binding, Molecular Mechanics Poisson– Boltzmann Surface Area (MM-PBSA) analysis was conducted using gmx_MMPBSA (latest version). Representative snapshots from the equilibrated 80–100 ns portion of the MD trajectory were extracted for energy calculations. The total binding free energy (ΔG□bind□) was calculated as the sum of van der Waals, electrostatic, polar solvation, and non-polar solvation energy components. Negative ΔG□ bind□ values were interpreted as indicative of spontaneous and favorable complex formation. These calculations provided quantitative support for the structural stability and binding affinity observed during simulation.

### Data analysis, visualization, and data presentation

All statistical analyses, including the Student’s two-tailed *t*-test and analysis of variance (ANOVA) with Tukey’s test, were performed using “ggplot2”, “ggpubr”, “dplyr”, “agricolae”, and “tibble” in RStudio (R 4.4.3). Biological replicates were indicated in the figure legends. All structural visualizations, trajectory inspections, and interaction maps were generated using Schrödinger Maestro (academic version, 2024-4) and Discovery Studio Visualizer (academic version). These platforms were employed to identify hydrogen-bond networks, hydrophobic interactions, and conformational changes across the simulation. Graphical outputs, including RMSD, RMSF, PCA, and DCCM plots, were generated in Python with Matplotlib and Seaborn to produce publication-quality figures. All numerical results were exported as CSV files, and textual summaries were compiled to provide a comprehensive view of system stability, residue flexibility, and binding interactions throughout the simulation workflow.

## RESOURCE AVAILABILITY

### Lead contact

Further information and requests for resources and reagents should be directed to and will be fulfilled by the lead contact, Byung-Wook Yun.

### Materials availability

Materials generated in this study are available upon request. For further details, contact the lead contact.

### Data and code availability

- All data reported in this article could be shared by the lead contact upon request.
- This article does not report original code.
- Accession numbers are listed in the key resources table.

## ACKNOWLEDGMENTS

We are grateful to the Global Korea Scholarship (GKS) of the Government of the Republic of Korea for supporting Ashim Kumar Das. Our graphical abstract was created with BioRender.com. Funding: This research was supported by the Basic Science Research Program through the National Research Foundation of Korea (NRF), funded by the Ministry of Education (RS-2023-00245922). This work was also supported by Korea Basic Science Institute (National Research Facilities and Equipment Center, 2021R1A6C101A416) funded by the Ministry of Education, biological materials specialized graduate program through the Korea Environmental Industry & Technology Institute (KEITI), funded by the Ministry of Climate, Energy and Environment (MCEE), and the Regional Innovation System & Education (RISE) Glocal 30 program through the Daegu RISE Center, funded by the Ministry of Education (MOE) and the Daegu, Republic of Korea (2025-RISE-03-001).

## AUTHOR CONTRIBUTIONS

A.K.D., B.-W.Y., and H.M. conceived and designed the research. A.K.D., D.-S.L., and S.-M.K. conducted plant experiments. H.M., A.K.D., and M.H. contributed to the structural analysis and interpretation of data. A.K.D. and H.M. wrote the 1^st^ draft. M.G.M. wrote and edited the manuscript. B.-W.Y. supervised and supported with resources and funding.

## DECLARATION OF INTERESTS

The authors declare no competing interests.

## STAR★METHODS

Detailed methods are provided in the online version of this paper and include the following:

- KEY RESOURCES TABLE
- METHOD DETAILS

- Plant materials and experimental design
- Assessment of photosynthetic pigments
- Quantification of H_2_O_2_ and GSH levels
- RNA isolation and qRT-PCR analysis
- Iodine staining and measurement of total soluble sugars and total carbohydrates
- Multiple sequence alignment and prediction of S-nitrosylation site
- Structure prediction using AlphaFold 3
- Structural modification for S-nitrosylation
- Small-molecule docking
- Peptide docking
- Protein and ligand preparation
- Molecular dynamics simulation
- Post-simulation analysis
- QUANTIFICATION AND STATISTICAL ANALYSIS

- Data analysis, visualization, and data presentation

## Notes

### Competing Interest Statement

The authors have declared no competing interest.

