## Supplementary File for "Integrative genetic and structural analyses reveal nitric oxide-driven regulation of ERFVII stability in Arabidopsis under low O_2_"

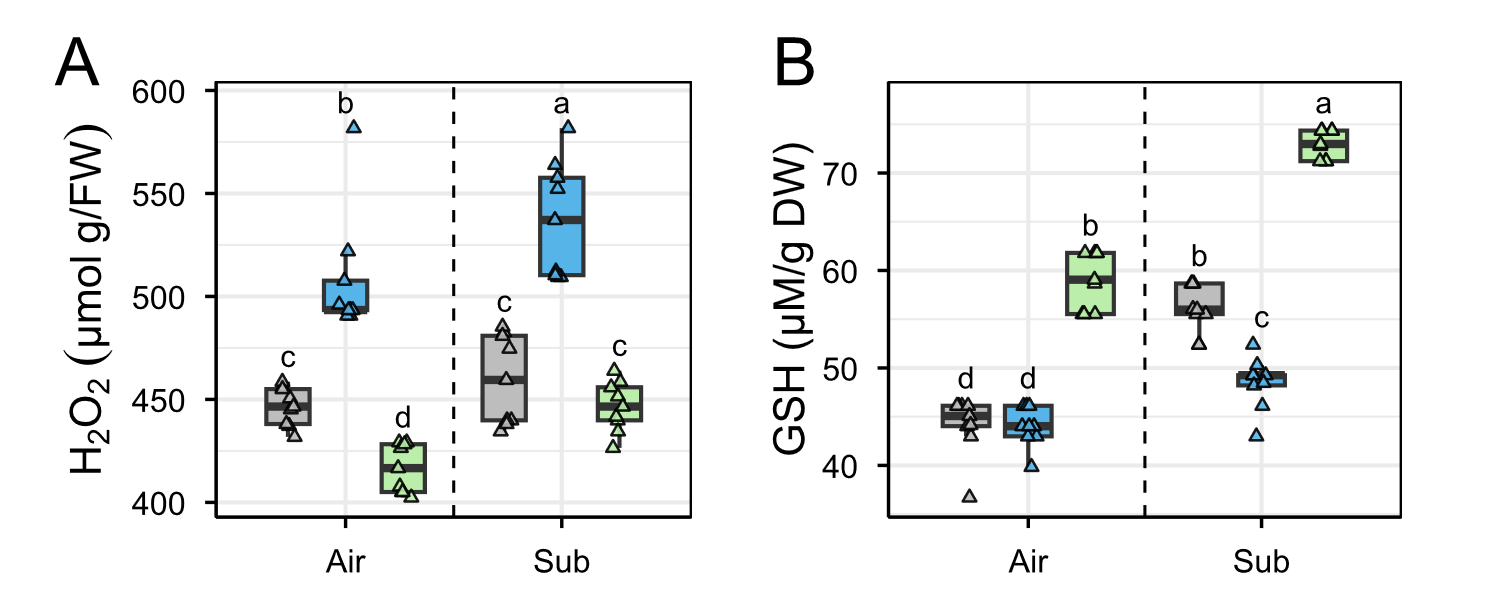

**Figure S1.** (F) H_2_O_2_ and (G) GSH levels in Col-0, *gsnor1-3*, and *gsnor1-1* plants exposed to air or 2 days of dark submergence (means ± SE, *n* = 4 biological replicates for EL, *n* = 9 technical replicates from 3 biological replicates). Different letters indicate statistically significant differences by two-way ANOVA and Tukey's HSD (*P* < 0.05).

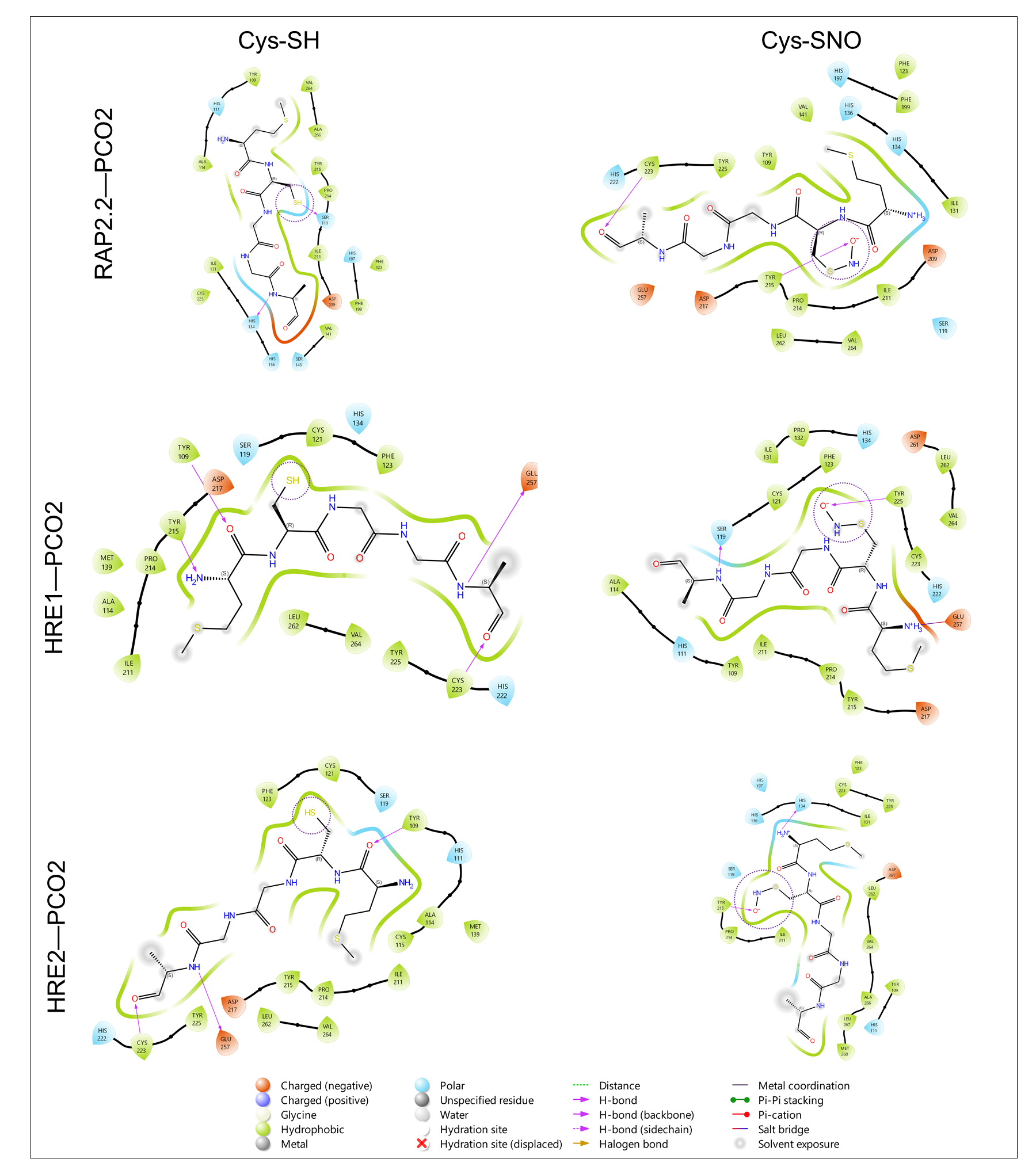

**Figure S2.** Heatmap representation of docking scores between proteins and peptides under both redox states. The 2D peptides-protein interaction of RAP2.2 (Peptides)-PCO2, HRE1 (Peptides)-PCO2, and (D, E) HRE2 (Peptides)-PCO2 are shown with or without redox modifications using Schrödinger Suite 2024-4 (academic version).

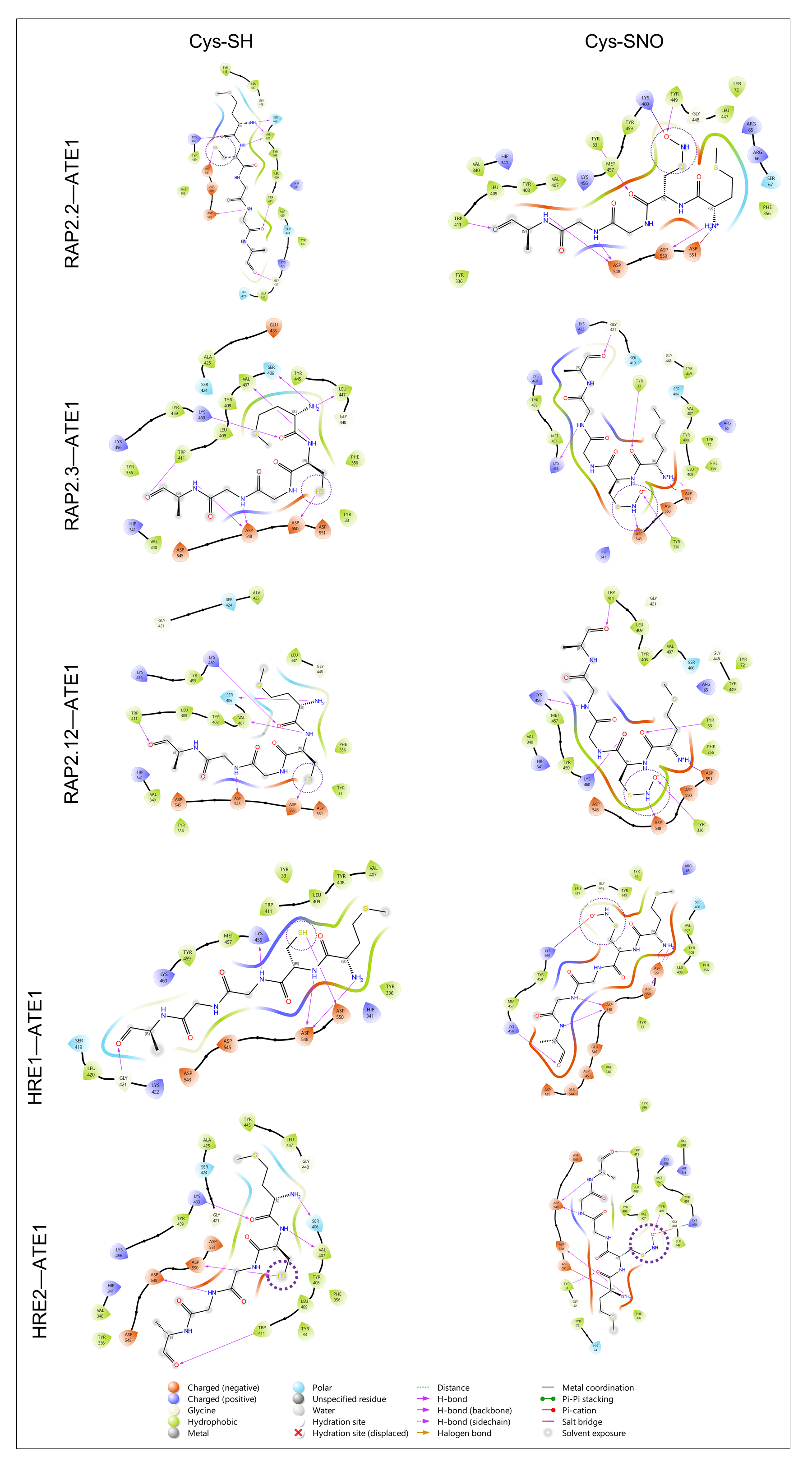

**Figure S3.** Heatmap representation of docking scores between proteins and peptides under both redox states. The 2D peptides-protein interaction of RAP2.2 (Peptides)-ATE1, RAP2.3 (Peptides)-ATE1, (D, E) RAP2.12 (Peptides)-ATE1, HRE1 (Peptides)-ATE1, and (D, E) HRE2 (Peptides)-ATE1 are shown with or without redox modifications using Schrödinger Suite 2024-4 (academic version).

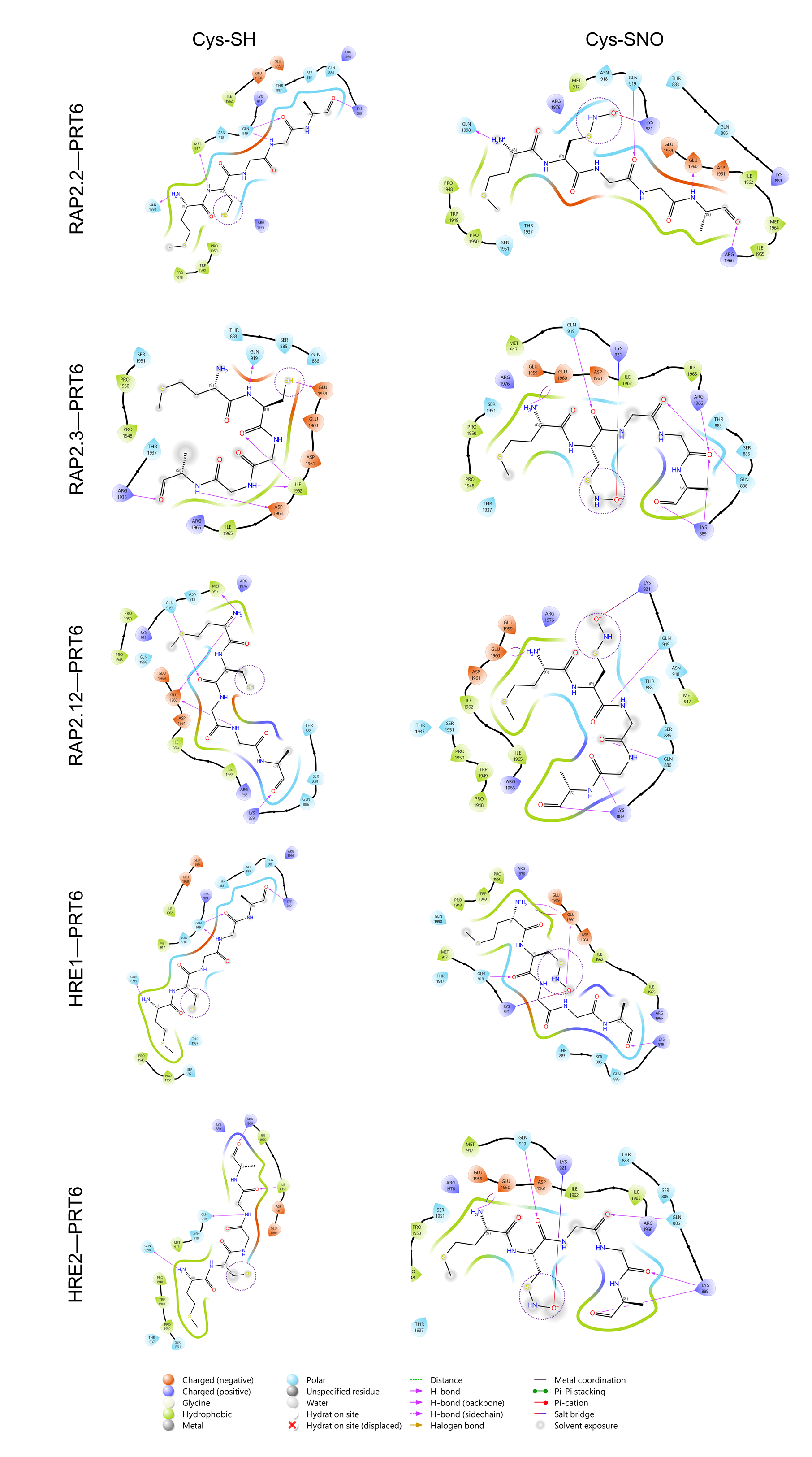

**Figure S4.** Heatmap representation of docking scores between proteins and peptides under both redox states. The 2D peptides-protein interaction of RAP2.2 (Peptides)-PRT6, RAP2.3 (Peptides)-PRT6, (D, E) RAP2.12 (Peptides)-PRT6, HRE1 (Peptides)-PRT6, and (D, E) HRE2 (Peptides)-PRT6 are shown with or without redox modifications using Schrödinger Suite 2024-4 (academic version).

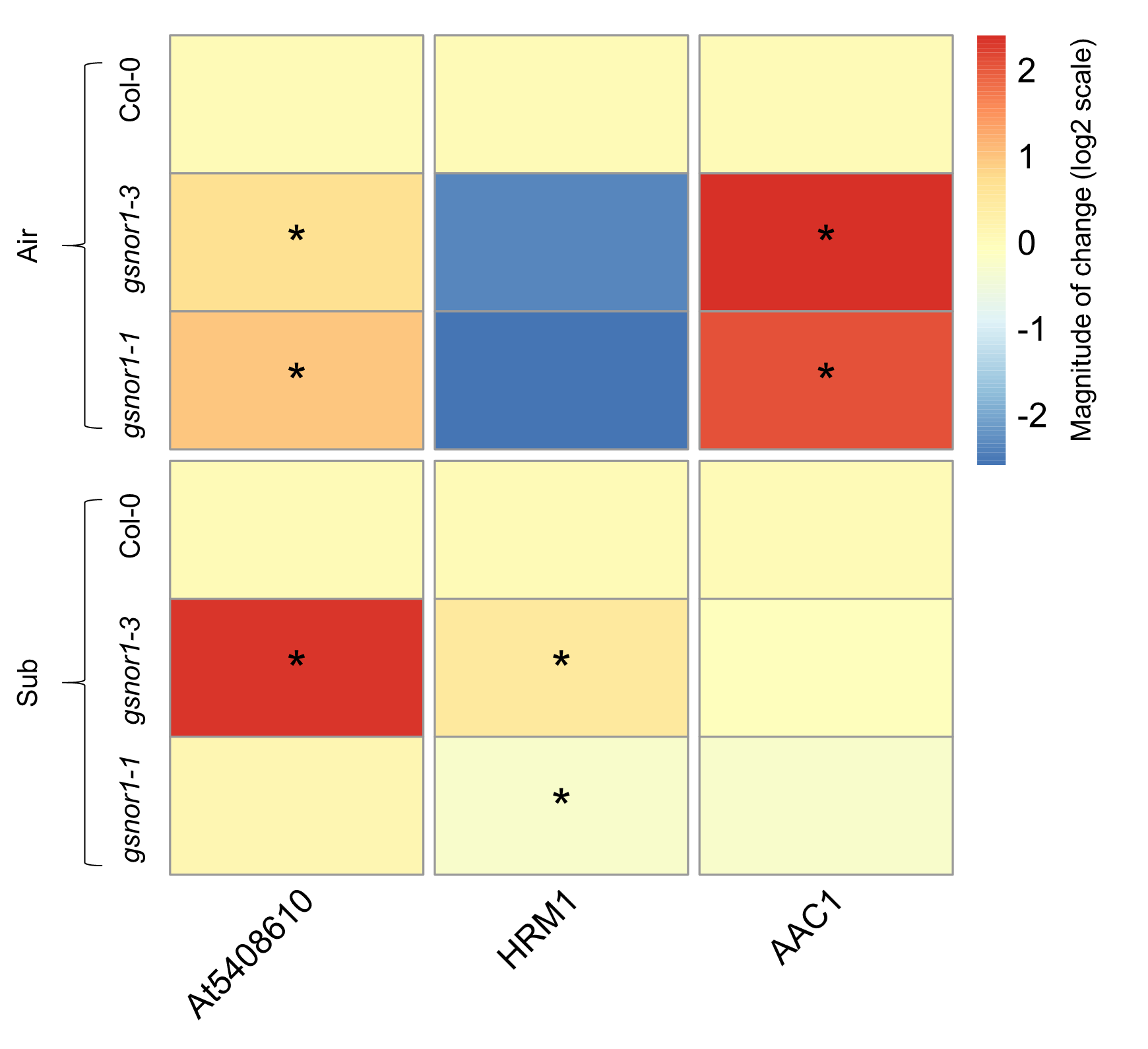

**Figure S5.** Expression pattern of genes involved in energy metabolism. Heatmap representation of relative gene expression, assessed by qualitative real-time polymerase chain reaction (qRT-PCR), in Col-0, *gsnor1-3*, and *gsnor1-1* after 24 h of dark submergence. Means (±SE) are calculated from 3 biological replicates, and the heatmap is displayed on a log2 scale. Asterisks indicate statistically significant differences by two-tailed Student *t*-test (*p < 0.05, **p< 0.01, ***p< 0.01).

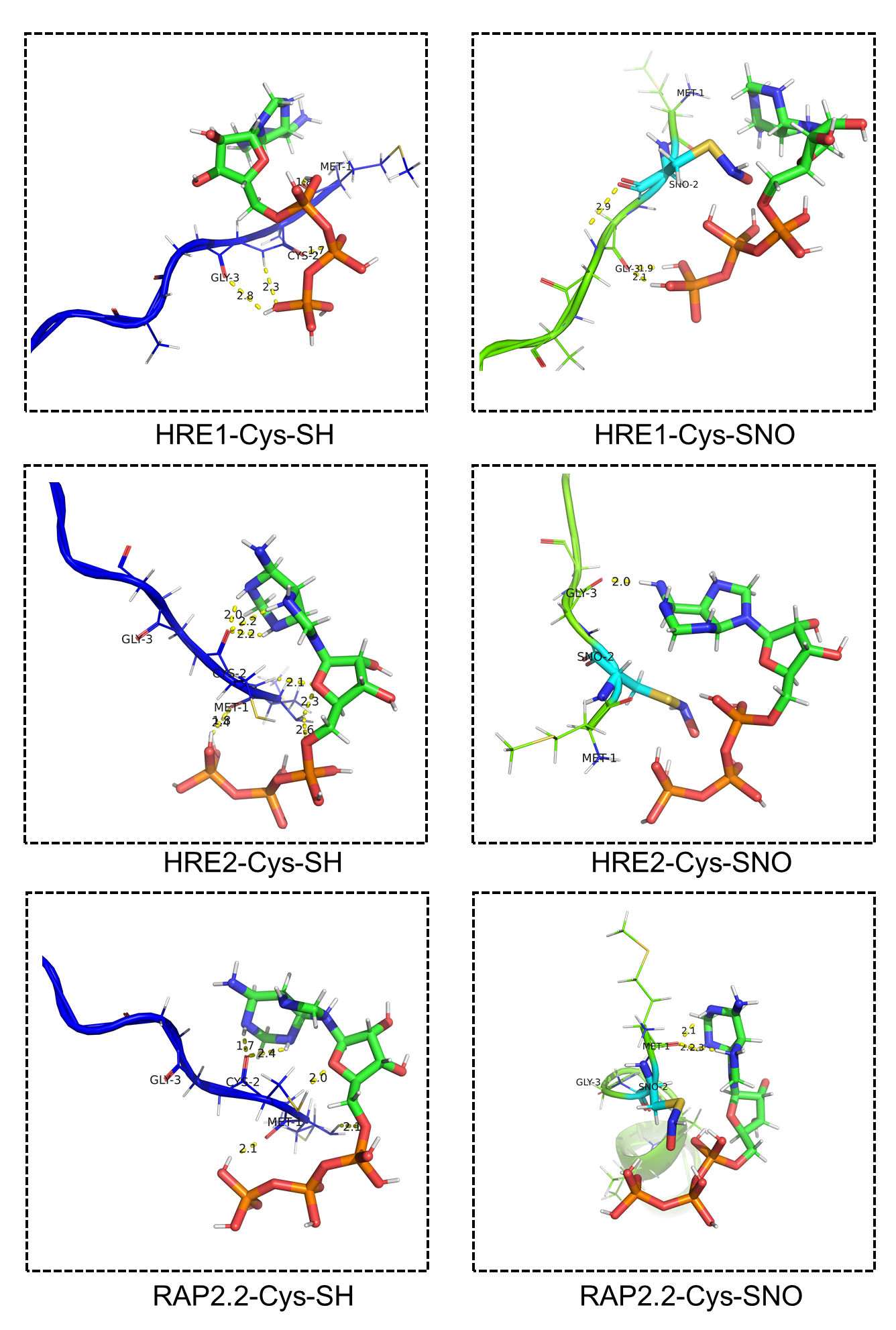

**Figure S6.** The 3D protein-ligand interaction of HRE1-ATP, HRE2-ATP, and RAP2.2-ATP is shown with or without redox modifications using PyMOL 3.1.6.1 (academic version).

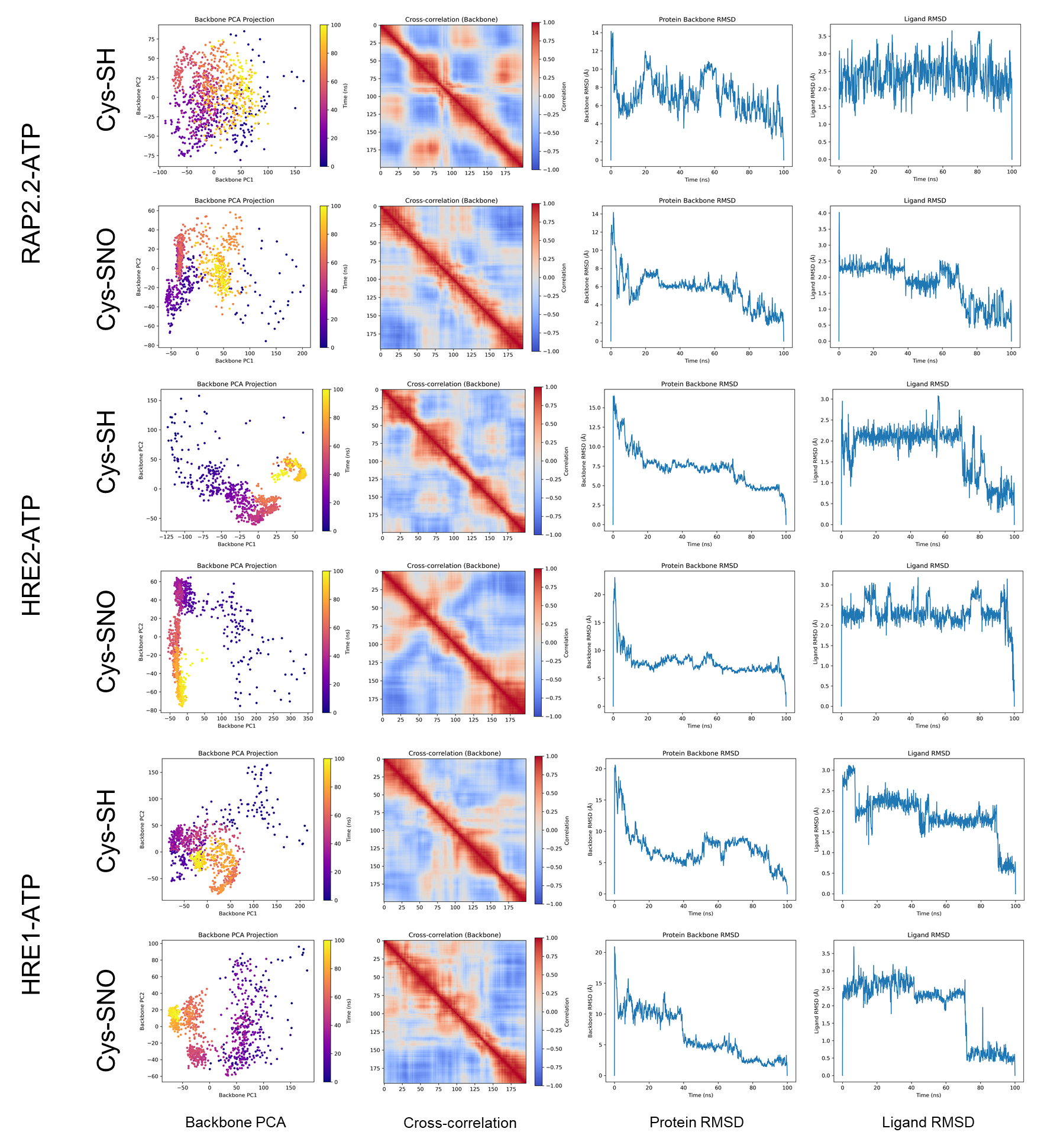

**Figure S7.** Molecular dynamics simulation analysis of RAP2.2-ATP, HRE2-ATP, and HRE1-ATP under reduced (Cys-SH) and S-nitrosylated (Cys-SNO) redox states. Backbone PCA projection showing conformational sampling, colored by time (purple = early, yellow = late). Dynamic cross-correlation matrices of backbone motions; red = correlated, blue = anti-correlated residue pairs. Protein backbone RMSD over time, reflecting structural stability relative to the initial docked conformation. Ligand (ATP) RMSD over time, indicating binding pocket stability throughout the simulation.

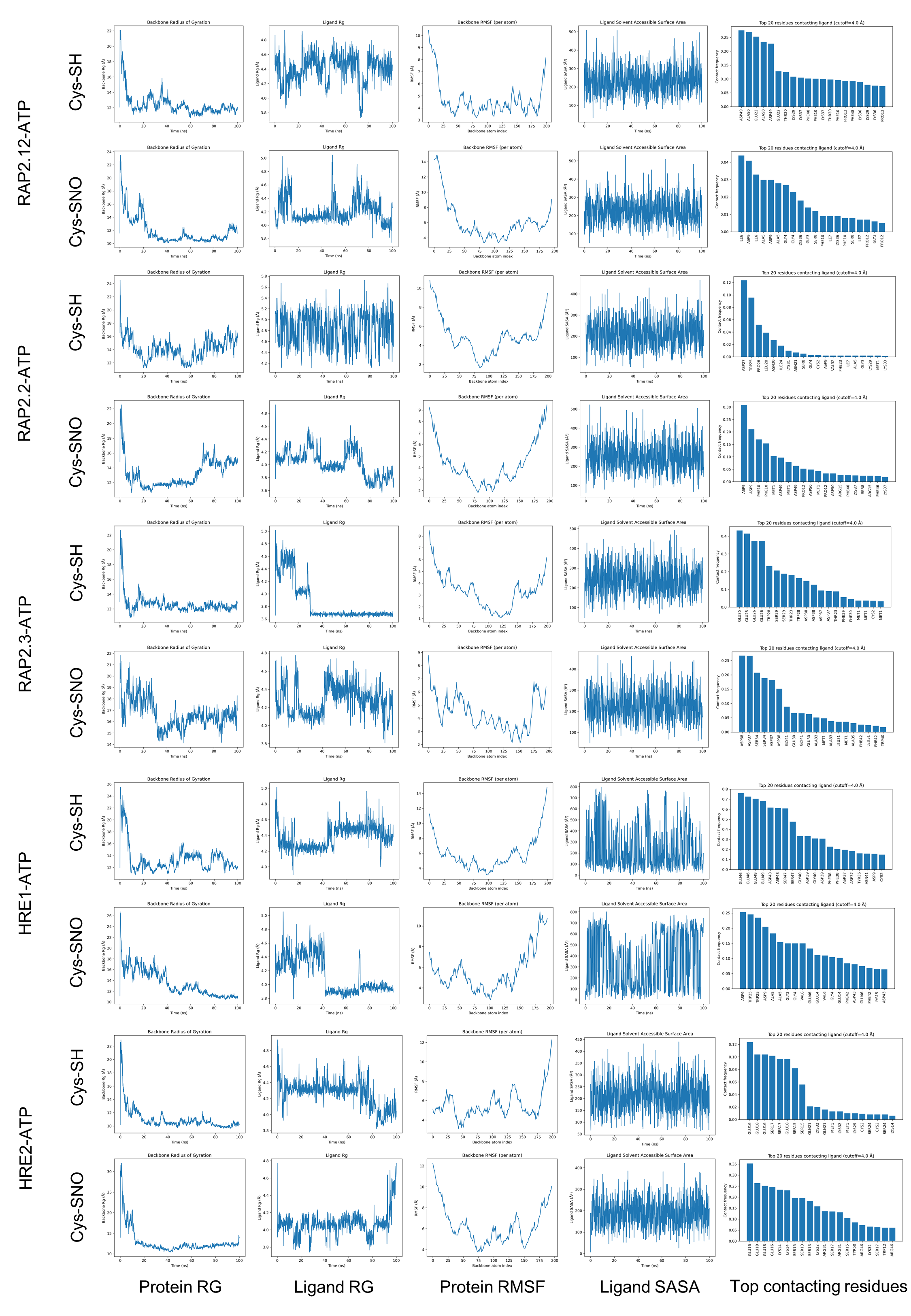

**Figure S8.** Molecular dynamics simulation analysis of ERVIIs-ATP under reduced (Cys-SH) and S-nitrosylated (Cys-SNO) redox states. Protein RG, ligand RG, protein RMSF, ligand SASA, and top contacting residues analysis between Cys-SH and Cys-SNO of group-VII ethylene response factor (ERFVII). RG, radius of gyration; RMSF, root mean square fluctuation; SASA, solvent accessible surface area.

**Table S1.** Gene-specific primer lists.

| **For qRT-PCR** | Sequence (5' --> 3') |
| --- | --- |
| *AtUBQ5 F* | GGAATCGACGCTTCATCTCG |
| *AtUBQ5 R* | ACTCCTTCCTCAAACGCTGA |
| *AtRAP2.12 F* | AGCGAGTTTATTTGGCCGGA |
| *AtRAP2.12 R* | AAAACTGAACCTTCCGCAGC |
| *AtRAP2.2 F* | CGCTGAGCAAGCTGAGAAAT |
| *AtRAP2.2 R* | TCCTCAGCAGTGTCGAATGT |
| *AtRAP2.3 F* | CCGATTATGCCCCTCTCGTC |
| *AtRAP2.3 R* | GGGATGGAGTTTGGAGGTGG |
| *AtHRE1 F* | ACATAGCGCCGGAGAAGATT |
| *AtHRE1 R* | CCATCTGATGCTGAGCCTGAA |
| *AtPCO1 F* | GGTTTAACTCCGACCATGCC |
| *AtPCO1 R* | AACACCAGAAGGTGGCAAAC |
| *AtPCO2 F* | TTCGCTGATGGCAAATCTGG |
| *AtPCO2 R* | GATCGTCCGGTCACTGTAGA |
| *AtATE1 F* | AGCCTATCGTCCATCTGAGC |
| *AtATE1 R* | GCTGGTTCCACGAGAGTTTC |
| *AtPRT6 F* | AATCGAGCATCCGATCCAGT |
| *AtPRT6 R* | TGGGCGACATGCACAATAAG |
| *AT5G08670 F* | ATTTCCGTGATGCTGAAGGC |
| *AT5G08670 R* | AAGCCAGAGTTGGCTGGTAT |
| *AtAAC1 F* | CGTCTGGTGAAGCTGTCAAG |
| *AtAAC1 R* | CCGAAGACAATCAGCTGCAA |
| *AtHRM1 F* | GCAGCCCAGTCGTCTACTAA |
| *AtHRM1 R* | AGAAAGAGCTAACGGCGAGA |
| **For genotyping** | Sequence (5' --> 3') |
| *Atgsnor1-1_F* | TATATAATGGTTCGACGCATATTT |
| *Atgsnor1-1_R* | GTCGGGTCGGGTCGTTAATA |
| *Atgsnor1-1_Sail LB* | TTCATAACCAATCTGGATACA |
| *Atgsnor1-3_F* | TATATAATGGTTCGACGCATA |
| *Atgsnor1-3_R* | CCACCAACACTCTCAACAATC |
| *Atgsnor1-3_GABI LB* | ATATTGACCATCATACTCATT |
